# Resolving the fibrotic niche of human liver cirrhosis using single-cell transcriptomics

**DOI:** 10.1101/766113

**Authors:** P Ramachandran, R Dobie, JR Wilson-Kanamori, EF Dora, BEP Henderson, RS Taylor, KP Matchett, JR Portman, M Efremova, R Vento-Tormo, NT Luu, CJ Weston, PN Newsome, EM Harrison, DJ Mole, SJ Wigmore, JP Iredale, F Tacke, JW Pollard, CP Ponting, JC Marioni, SA Teichmann, NC Henderson

**Author notes:** Address correspondence to: Prakash Ramachandran, University of Edinburgh Centre for Inflammation Research, The Queen’s Medical Research Institute, Edinburgh BioQuarter, 47 Little France Crescent, Edinburgh, UK., EH16 4TJ.; or Neil Henderson, University of Edinburgh Centre for Inflammation Research, The Queen’s Medical Research Institute, Edinburgh BioQuarter, 47 Little France Crescent, Edinburgh, UK., EH16 4TJ.

## Abstract

Currently there are no effective antifibrotic therapies for liver cirrhosis, a major killer worldwide. To obtain a cellular resolution of directly-relevant pathogenesis and to inform therapeutic design, we profile the transcriptomes of over 100,000 primary human single cells, yielding molecular definitions for the major non-parenchymal cell types present in healthy and cirrhotic human liver. We uncover a novel scar-associated TREM2^+^CD9^+^ macrophage subpopulation with a fibrogenic phenotype, that has a distinct differentiation trajectory from circulating monocytes. In the endothelial compartment, we show that newly-defined ACKR1^+^ and PLVAP^+^ endothelial cells expand in cirrhosis and are topographically located in the fibrotic septae. Multi-lineage ligand-receptor modelling of specific interactions between the novel scar-associated macrophages, endothelial cells and collagen-producing myofibroblasts in the fibrotic niche, reveals intra-scar activity of several major pathways which promote hepatic fibrosis. Our work dissects unanticipated aspects of the cellular and molecular basis of human organ fibrosis at a single-cell level, and provides the conceptual framework required to discover rational therapeutic targets in liver cirrhosis.

## Main

Liver cirrhosis is a major global healthcare burden. Recent estimates suggest that 844 million people worldwide have chronic liver disease, with a mortality rate of two million deaths per year and a rising incidence^1^. In health, the liver serves a myriad of functions including detoxification, metabolism, bile production and immune surveillance. Chronic liver disease, the result of iterative liver injury secondary to any cause, results in progressive fibrosis, disrupted hepatic architecture, vascular changes and aberrant regeneration, defining characteristics of liver cirrhosis^2^. Importantly, the degree of liver fibrosis predicts adverse patient outcomes, including the development of cirrhosis-related complications, hepatocellular carcinoma and death^3^. Hence, there is a clear therapeutic imperative to develop effective anti-fibrotic approaches for patients with chronic liver disease^4–7^.

Liver fibrosis involves a complex, orchestrated interplay between multiple non-parenchymal cell (NPC) lineages including immune, endothelial and mesenchymal cells spatially located within areas of scarring, termed the fibrotic niche. Despite rapid progress in our understanding of the cellular interactions underlying liver fibrogenesis accrued using rodent models, there remains a significant ’translational gap’ between putative targets and effective patient therapies^4, 5^. This is in part due to the very limited definition of the functional heterogeneity and interactome of cell lineages that contribute to the fibrotic niche of human liver cirrhosis, which is imperfectly recapitulated by rodent models^4, 6^.

Single-cell RNA sequencing (scRNA-seq) has the potential to deliver a step change in both our understanding of healthy tissue homeostasis as well as disease pathogenesis, allowing the interrogation of individual cell populations at unprecedented resolution^8–10^. Here, we have studied the mechanisms regulating human liver cirrhosis, using scRNA-seq to analyse the transcriptomes of 106,616 single cells obtained from ten healthy and cirrhotic human livers and peripheral blood.

Our data define: (1) a single-cell atlas of non-parenchymal cells in healthy and cirrhotic human liver; (2) a new subpopulation of scar-associated TREM2^+^CD9^+^ pro-fibrotic macrophages; (3) new subpopulations of scar-associated ACKR1^+^ and PLVAP^+^ endothelial cells; and (4) key ligand-receptor interactions between novel scar-associated macrophages, endothelial subpopulations and collagen-producing myofibroblasts in the fibrotic niche. Thus, we have simultaneously identified a series of intra-scar fibrogenic pathways which represent hitherto unsuspected therapeutic targets for the treatment of liver fibrosis, whilst demonstrating the applicability of scRNA-seq to define pathogenic mechanisms for other human fibrotic disorders.

## Results

### Single cell atlas of human liver non-parenchymal cells

Hepatic NPC were isolated from fresh healthy and cirrhotic human liver tissue spanning a range of aetiologies of cirrhosis (Fig. 1a, Extended Data Fig. 1a). Importantly, to minimise artefacts^11^, we developed a rapid tissue processing pipeline, obtaining fresh non-ischaemic liver tissue taken by wedge biopsy prior to the interruption of the hepatic vascular inflow during liver surgery or transplantation, and immediately processing this for FACS. This enabled a workflow time of under three hours from patient to single-cell droplet encapsulation (Methods).

**Figure 1:**
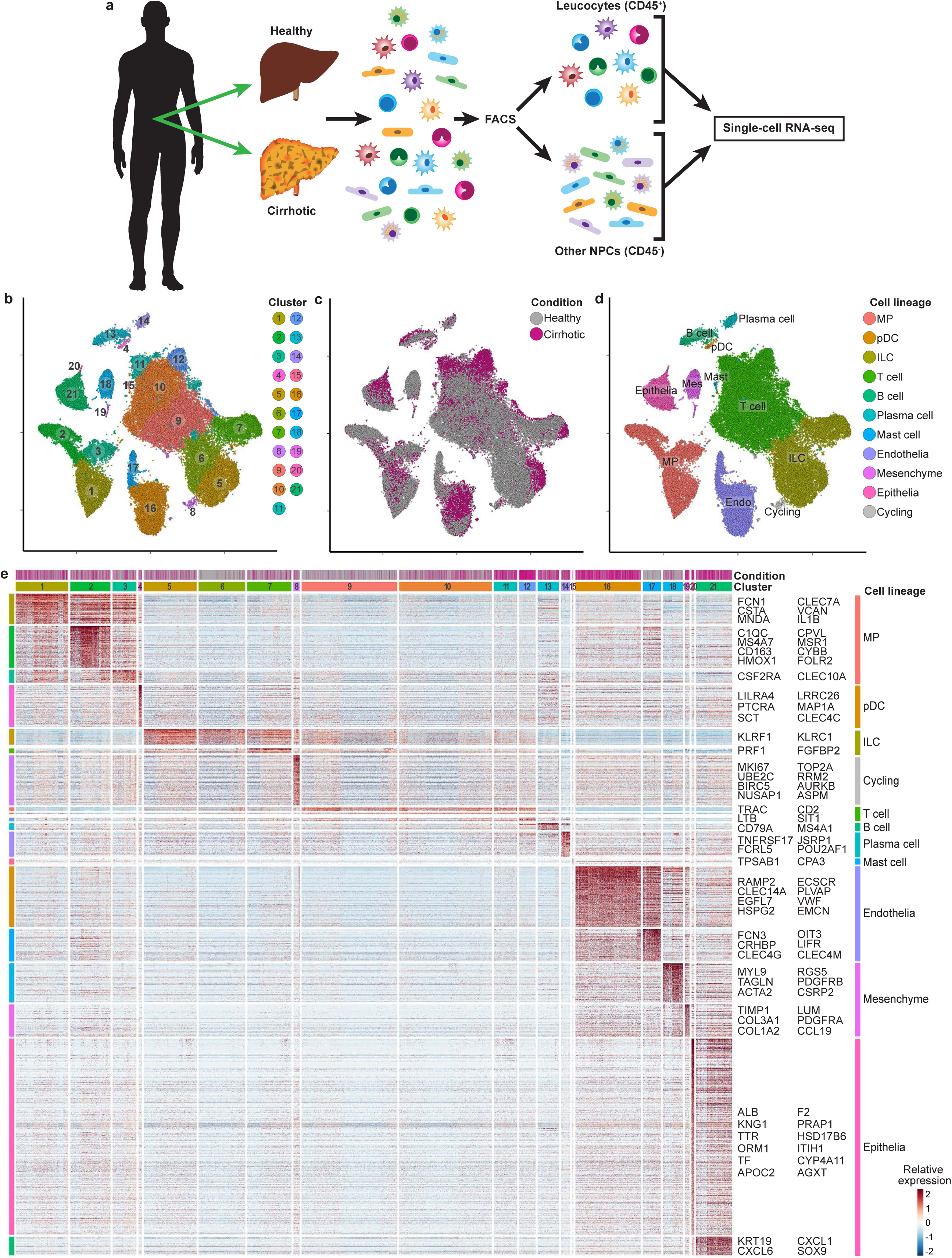
Single cell atlas of human liver non-parenchymal cells. **a**, Overview: extraction of non-parenchymal cells (NPC) from healthy or cirrhotic human liver, followed by fluorescence-activated cell sorting (FACS) into leucocyte (CD45^+^) and other NPC fractions (CD45^-^) for droplet-based 3’ single-cell RNA-seq. **b**, t-SNE visualisation: clustering (colour) 66,135 non-parenchymal cells (points; n=5 healthy and n=5 cirrhotic human livers). **c**, t-SNE visualisation: injury condition (colour; healthy versus cirrhotic). **d**, t-SNE visualisation: cell lineage (colour) inferred from expression of known marker gene signatures. Endo, endothelial cell; ILC, innate lymphoid cell; Mast, mast cell; Mes, mesenchymal cell; MP, mononuclear phagocyte; pDC, plasmacytoid dendritic cell. e, Scaled heatmap (red, high; blue, low): cluster marker genes (top, colour coded and numbered by cluster and colour coded by condition) and exemplar genes and lineage annotation labelled (right). Cells columns, genes rows.

We used an unbiased approach, FACS sorting viable single cells from liver tissue into broad leucocyte (CD45^+^) or other NPC (CD45^-^) fractions (Extended Data Fig. 1b), prior to scRNA-seq. To facilitate discrimination between liver-resident and circulating leucocytes, we also performed scRNA-seq on CD45^+^CD66b^-^ peripheral blood mononuclear cells (PBMCs) (Extended Data Fig. 1c, f). In total, we analysed 67,494 human cells from healthy (n=5) and cirrhotic (n=5) livers, 30,741 PBMCs from cirrhotic patients (n=4) and compared our data with a publicly-available reference dataset of 8,381 PBMCs from a healthy donor.

Tissue cells and PBMCs could be partitioned into 21 distinct clusters, which we visualized using *t*-distributed stochastic neighbourhood embedding (t-SNE) (Extended Data Fig. 1d). Clusters were annotated using signatures and integrating with known lineage markers (Extended Data Fig. 1e; signature gene lists available in Supplementary Table 1). All PBMC datasets contained the major blood lineages, with excellent reproducibility between samples (Extended Data Fig. 1g, h). To generate an atlas of liver-resident cells, contaminating circulating cells were removed from the liver tissue datasets, by excluding individual cells from the tissue samples which mapped transcriptionally to blood-derived clusters 1 and 13 (Extended Data Fig. 1d).

Re-clustering the 66,135 liver-resident cells revealed 21 clusters (Fig. 1b), each containing cells from both healthy and cirrhotic livers (Fig. 1c). Gene signature analysis enabled annotation of each cluster by major cell lineage (Fig. 1d, Extended Data Fig. 2a, b). We noted heterogeneity in the post-normalised detected number of genes and unique molecular identifiers (UMIs) per cell dependent on cell lineage, with mononuclear phagocytes (MP) demonstrating increased transcriptional diversity (nGene; Healthy=1,513±10.5, Cirrhotic=1,912±10.8) and activity (nUMI; Healthy=5,499±58.4, Cirrhotic=7,386±66) in diseased livers (Extended Data Fig. 2c, d). All samples contained the expected cell lineages (Extended Data Fig. 2e, g) and reproducibility between livers was excellent for the main NPC populations (Extended Data Fig. 2f).

**Figure 2:**
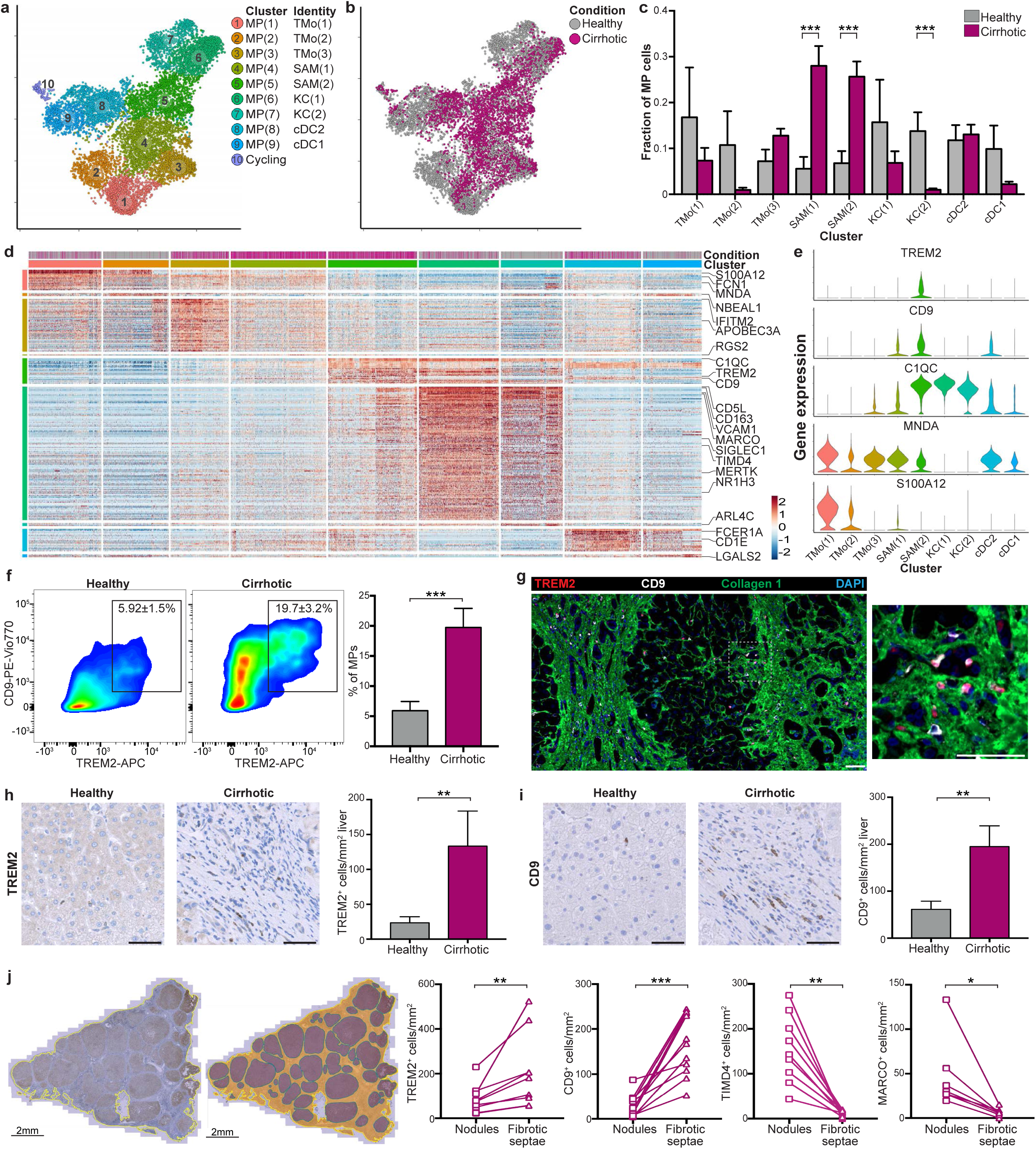
Identifying scar-associated macrophage subpopulations. **a**, t-SNE visualisation: clustering 10,737 mononuclear phagocytes (MP) into 10 clusters (colour and number). Annotation of clusters (identity). TMo, tissue monocyte; SAM, scar-associated macrophage; KC, Kupffer cell; cDC, conventional dendritic cell. **b**, t-SNE visualisation: annotating MP cells by injury condition (colour). **c**, Fractions of MP subpopulations in healthy (n=5) versus cirrhotic (n=5) livers, Wald. **d**, Scaled heatmap (red, high; blue, low): MP cluster marker genes (top, colour coded by cluster and condition), exemplar genes labelled (right). Cells columns, genes rows. **e**, Violin plots: scar-associated macrophage and tissue monocyte cluster markers. **f**, Representative flow cytometry plots: quantifying TREM2^+^CD9^+^ MP fraction by flow cytometry in healthy (n=2) versus cirrhotic (n=3) liver, Wald. **g**, Representative immunofluorescence micrograph, cirrhotic liver: TREM2 (red), CD9 (white), collagen 1 (green), DAPI (blue). Scale bar, 50μm. **h**, Automated cell counting: TREM2 staining, healthy (n=10) versus cirrhotic (n=9) liver, Mann-Whitney. **i**, Automated cell counting: CD9 staining, healthy (n=12) versus cirrhotic (n=10) liver, Mann-Whitney. **j**, Topographically assessing scar-associated macrophages: exemplar tissue segmentation (left), stained section morphologically segmented into fibrotic septae (orange) and parenchymal nodules (purple)). TREM2^+^, CD9^+^, TIMD4^+^ and MARCO^+^ automated cell counts (right) in parenchymal nodules versus fibrotic septae, Wilcoxon. Error bars, s.e.m.; * p-value<0.05; ** p-value < 0.01; *** p-value < 0.001.

We used an area-under-curve classifier to identify cell subpopulation markers across all 21 clusters and 11 lineages (Fig. 1e; Supplementary Tables 2 and 3). Expression of collagens type I and type III, the main extracellular matrix components of the fibrotic niche, was restricted to cells of the mesenchymal lineage (Fig. 1e). To gain further resolution on NPC heterogeneity, we then iterated clustering and marker gene identification on each lineage in turn, for example defining 11 clusters of T cells and innate lymphoid cells (ILCs) (Extended Data Fig. 3a) and four clusters of B cells and plasma cells (Extended Data Fig. 3f, g, Supplementary Table 6). No major differences in B cell or plasma cell composition between healthy and cirrhotic livers were observed (Extended Data Fig. 3h), and plasmacytoid dendritic cells (pDC) showed no additional heterogeneity.

**Figure 3:**
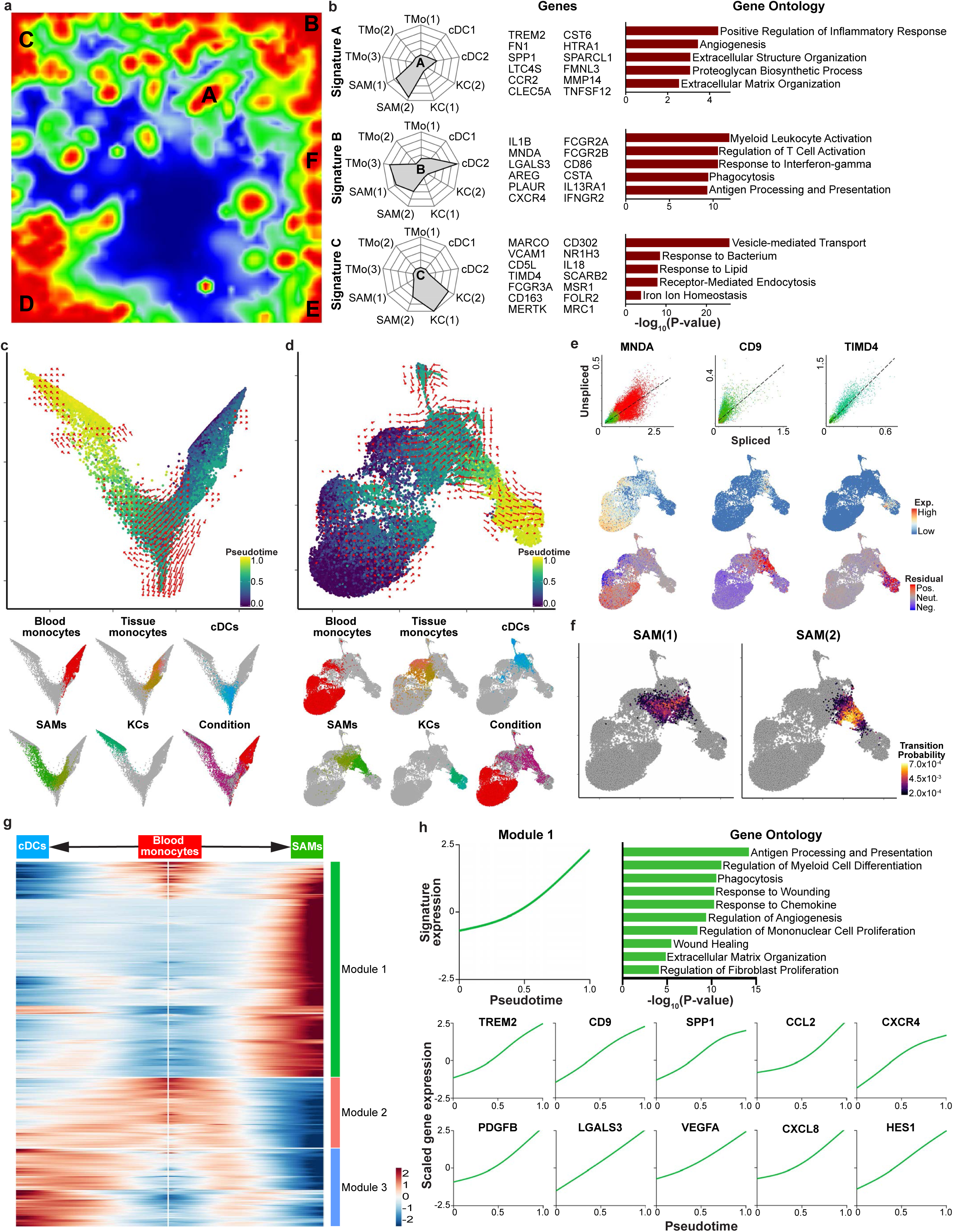
Fibrogenic phenotype of scar-associated macrophages. **a**, Self-Organising Map (SOM; 60×60 grid): smoothed scaled metagene expression of mononuclear phagocyte (MP) lineage. 20,952 genes, 3,600 metagenes, 44 signatures. A-F label metagene signatures overexpressed in one or more MP subpopulations. **b**, Radar plots (left): metagene signatures A-C showing distribution of signature expression across MP subpopulations, exemplar genes (middle) and gene ontology (GO) enrichment (right). **c**, Diffusion map visualisation, blood monocytes and liver-resident MP lineages (23,075 cells), annotating *monocle* pseudotemporal dynamics (purple to yellow). RNA velocity field (red arrows) visualised using Gaussian smoothing on regular grid. Below: Annotation of MP subpopulation, injury condition. **d**, UMAP visualisation, blood monocytes and liver-resident MP lineages, annotating *monocle* pseudotemporal dynamics (purple to yellow). RNA velocity field (red arrows) visualised using Gaussian smoothing on regular grid. Below: Annotation of MP subpopulation, injury condition. **e**, Unspliced-spliced phase portraits (top row), cells coloured as in **d**, for monocyte (*MNDA*), SAM (*CD9*) and KC marker genes (*TIMD4*). Cells plotted above or below the steady-state (black dashed line) indicate increasing or decreasing expression of gene, respectively. Spliced expression profile for genes (middle row; red high, blue low). Unspliced residuals (bottom row), positive (red) indicating expected upregulation, negative (blue) indicating expected downregulation for genes. *MNDA* displays negative velocity in SAMs, *CD9* displays positive velocity in monocytes and SAMs, *TIMD4* velocity is restricted to KCs. **f**, UMAP visualisation, transition probabilities per SAM subpopulation, indicating for each cell the likelihood of transition into either SAM(1) or SAM(2), calculated using RNA velocity (yellow high; purple low; grey below threshold of 2×10^-4^). **g**, Scaled heatmap (red, high; blue low): cubic smoothing spline curves fitted to genes differentially expressed across blood monocyte-to-SAM (right arrow) and blood monocyte-to-cDC (left arrow) pseudotemporal trajectories, grouped by hierarchical clustering (k=3). Gene co-expression modules (colour) labelled right. **h**, Cubic smoothing spline curve fitted to averaged expression of all genes in module 1, along monocyte-SAM pseudotemporal trajectory, selected GO enrichment (right) and curves fit to exemplar genes (below).

To further annotate the 11 T cell and ILC clusters (36,900 cells from 10 livers) we assessed expression of known markers (Extended Data Fig. 3c) and computationally identified differential marker genes (Extended Data Fig. 3d, Supplementary Table 4). We also performed imputation of gene dropouts, which enhanced detection of discriminatory marker genes for each cluster but did not yield additional T cell or ILC subpopulations (Extended Data Fig. 3e, Supplementary Table 5). All T cell and ILC clusters expressed tissue residency markers *CD69* and *CXCR4*. Clusters 1 and 2 were CD4^+^ T cells, with CD4^+^ T cell(2) expanding significantly in cirrhotic livers (Extended Data Fig. 3a, b, e) and expressing *SELL* and *CCR7,* indicating an expansion of memory CD4^+^ T cells in liver cirrhosis. Sparse expression of *FOXP3, RORC*, *IL17A* and *IFNG* in both CD4^+^ T cell subpopulations suggested the presence of Tregs, Th17 and Th1 cells in these clusters. Clusters 3, 4 and 5 were CD8^+^ T cells, with features of effector T cells expressing *GZMA*, *GZMH* and *IFNG*. Two resident CD56^bright^ IL7R^-^ NK cell clusters were defined (NK cell(1) and NK cell(2)), as well as a distinct cytotoxic CD56^dim^ NK cell population (cNK), with specific expression of *FCGR3A* and *GZMB*. No expansion of these populations was observed in cirrhotic livers.

We provide an interactive gene browser freely-available online ((http://www.livercellatlas.mvm.ed.ac.uk), to allow assessment of individual gene expression both in all human liver NPC and in specific lineages, comparing healthy versus cirrhotic livers.

### Distinct macrophage subpopulations inhabit the fibrotic niche

Macrophages are critical to tissue homeostasis and wound-healing^12^. Previous studies have highlighted phenotypically-distinct macrophage populations orchestrating both liver fibrosis progression and regression in rodent models^13, 14^, with preliminary evidence of heterogeneity in fibrotic human livers^15^. Here, we define unique subpopulations of macrophages which populate the fibrotic niche of cirrhotic human livers. Unsupervised clustering of all 10,737 mononuclear phagocytes (1,074±153 cells from each liver), isolated from the combined liver-resident cell dataset, identified nine MP clusters and one cluster of proliferating MP cells (Fig. 2a). We annotated these nine clusters as subpopulations of scar-associated macrophages (SAM), Kupffer cells (KC), tissue monocytes (TMo), and conventional dendritic cells (cDC) (Fig. 2a; see below). Strikingly, clusters MP(4) and MP(5), named SAM(1) and SAM(2) respectively, were expanded in cirrhotic livers (Fig. 2b), a finding that was confirmed by quantification of the MP cell composition of each liver individually (Fig. 2c), and reproduced in all cirrhotic livers irrespective of liver disease aetiology.

To enable MP cell annotation, we initially assessed expression of known MP marker genes (Extended Data Fig. 4a), classifying clusters MP(8) and MP(9) as conventional dendritic cells, cDC2 and cDC1 respectively, based on *CD1C* and *CLEC9A* specificity. However, the remaining markers did not demarcate the other monocyte and macrophage subpopulations. Instead, these were identified using differential expression analysis across all MP clusters (Fig. 2d, Supplementary Table 7). Clusters MP(1), MP(2) and MP(3) were distinguished by expression of S100 genes, *FCN1, MNDA* and *LYZ*, in keeping with a tissue monocyte (TMo) phenotype and informing annotation as TMo(1), TMo(2) and TMo(3) respectively (Fig. 2d, e, Extended Data Fig. 4a, Supplementary Table 7).

**Figure 4:**
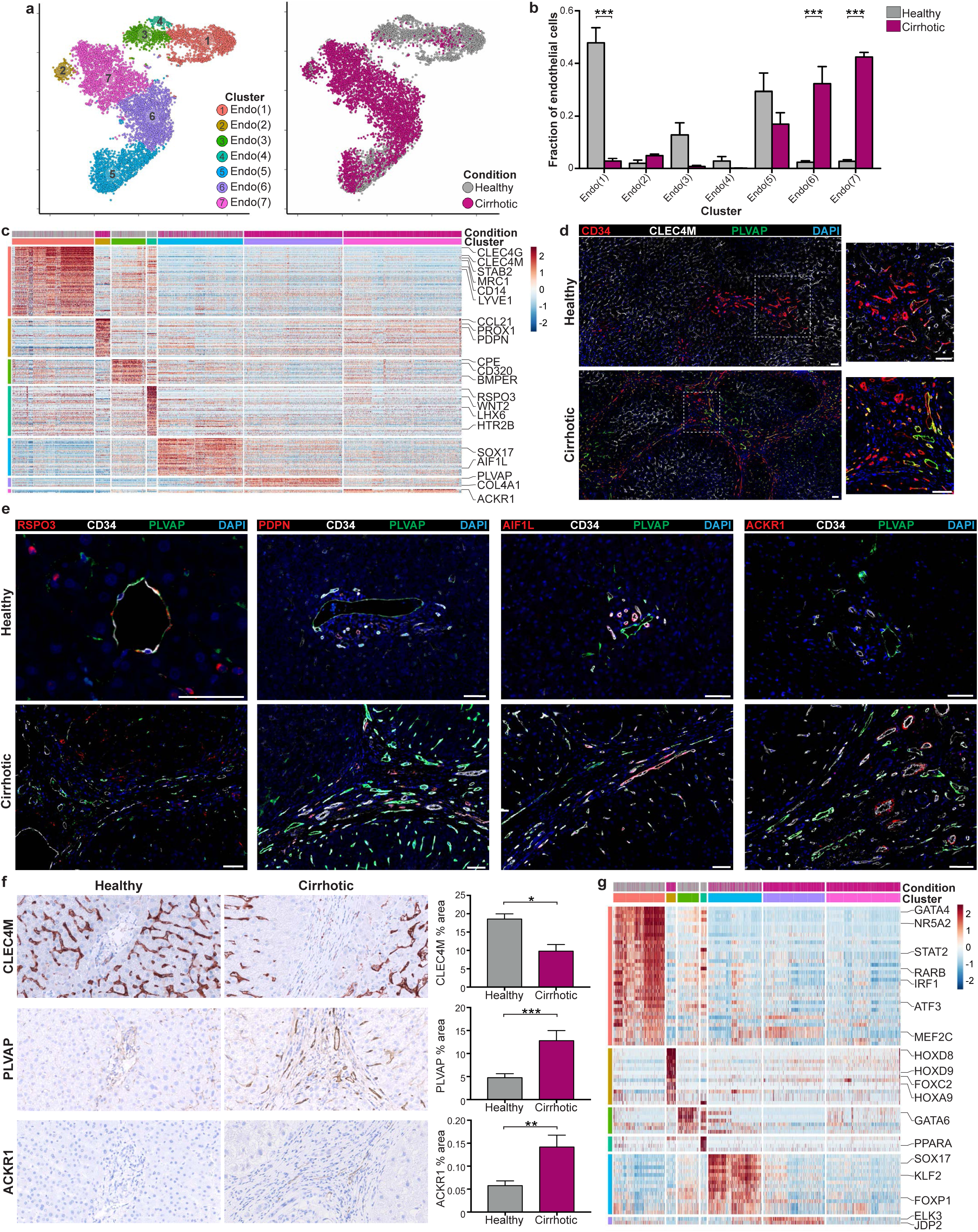
Identifying scar-associated endothelial subpopulations. **a**, t-SNE visualisation: clustering 8,020 endothelial cells, annotating injury condition. **b**, Fractions of endothelial subpopulations in healthy (n=4) versus cirrhotic (n=3) livers, Wald. **c**, Scaled heatmap (red, high; blue, low): endothelial cluster marker genes (colour coded top by cluster and condition), exemplar genes labelled right. Cells columns, genes rows. **d**, Representative immunofluorescence micrograph, healthy versus cirrhotic liver: CD34 (red), CLEC4M (white), PLVAP (green), DAPI (blue). **e**, Representative immunofluorescence micrographs, healthy versus cirrhotic liver: RSPO3, PDPN, AIF1L or ACKR1 (red), CD34 (white), PLVAP (green), DAPI (blue). **f**, Digital morphometric pixel quantification: CLEC4M staining healthy (n=5) versus cirrhotic (n=8), PLVAP staining healthy (n=11) versus cirrhotic (n=11), ACKR1 staining healthy (n=10) versus cirrhotic (n=10), Mann-Whitney. **g**, Scaled heatmap (red, high; blue, low): endothelial cluster marker transcription factor regulons (colour coded top by cluster and condition), exemplar regulons labelled right. Cells in columns, regulons in rows. Scale bars, 50μm. Error bars, s.e.m.; * p-value < 0.05, ** p-value < 0.01, *** p-value < 0.001.

Clusters MP(6) and MP(7) were enriched in *CD163*, *MARCO*, *TIMD4* and *CD5L* (Extended Data Fig. 4b); multiplex immunofluorescence staining confirmed these as Kupffer cells (KC; resident liver macrophages), facilitating annotation of these clusters as KC(1) and KC(2) respectively (Extended Data Fig. 4c). Application of these markers enabled the definitive distinction between KCs and other MP cells for the first time in human liver tissue. KCs displayed characteristic morphology and sinusoidal topography in healthy livers but were absent from areas of scarring in cirrhotic livers (Extended Data Fig. 4c). A lack of *TIMD4* expression distinguished KC(2) from KC(1) (Extended Data Fig. 4b); CD163^+^MARCO^+^TIMD4^-^ cells were identifiable in healthy livers but rare in cirrhotic livers (Extended Data Fig. 4c), concordant with a significant reduction of KC(2) cells in cirrhosis (Fig. 2c). Automated histological cell counting demonstrated TIMD4^+^ cell numbers to be equivalent between healthy and cirrhotic livers, but showed a loss of MARCO^+^ cells, consistent with selective reduction in MARCO^+^TIMD4^-^ KCs in liver fibrosis (Extended Data Fig. 4d, e).

Scar-associated clusters SAM(1) and SAM(2), expanded in diseased livers and expressed the unique markers *TREM2* and *CD9* (Fig. 2d, e). These newly-defined macrophages displayed a hybrid phenotype, with features of both tissue monocytes and KCs (Fig. 2d, e), analogous to monocyte-derived macrophages in murine liver injury models^14, 16^. Multi-colour flow cytometry confirmed expansion of these TREM2^+^CD9^+^ macrophages in human fibrotic livers (Fig. 2f, Extended Data Fig. 4f). Tissue immunofluorescence staining and single-molecule fluorescent in situ hybridization (smFISH) demonstrated the presence of TREM2^+^MNDA^+^ and CD9^+^MNDA^+^ macrophages in fibrotic livers (Extended Data Fig. 4g-i). Multiplex immunofluorescence further confirmed the presence of TREM2^+^CD9^+^ cells localised in collagen-positive scar regions in cirrhotic livers (Fig. 2g), and automated cell counting of stained sections confirmed expansion of TREM2^+^ and CD9^+^ cells in cirrhotic livers (Fig. 2h, i).

Strikingly, TREM2^+^ and CD9^+^ cells were rarely identified in the parenchyma of healthy livers, but were consistently located within areas of scar in cirrhotic livers. To confirm this, automated cell counting was applied to immunohistochemically-stained cirrhotic livers morphologically segmented into regions of fibrotic septae and parenchymal nodules (Fig. 2j). This demonstrated a significant accumulation of TREM2^+^ and CD9^+^ cells in fibrotic regions, whilst negligible numbers of KCs populated the fibrotic septae (Fig. 2j). Hence, we annotated TREM2^+^CD9^+^ MP cells as scar-associated macrophages.

Local proliferation has been shown to play a significant role in the expansion of macrophage subpopulations at sites of inflammation and fibrosis in experimental rodent models^14, 17, 18^, but has not been extensively characterised in human inflammatory disorders. To investigate MP proliferation in human liver fibrosis, we isolated the cycling MP cluster (Fig. 2a; cluster 10), which was enriched for multiple cell cycle-related genes (Supplementary Table 7). Cycling MP cells subclustered into four, yielding cDC1, cDC2, KCs and scar-associated macrophage subpopulations (Extended Data Fig. 4j). We observed a significant expansion of cycling SAMs in cirrhosis, representing 1.70±0.52% of total TREM2^+^ MP cells in cirrhotic livers (Extended Data Fig. 4k). In contrast 0.99±0.63% of KCs were proliferating in healthy livers, with none detected in cirrhotic livers (Extended Data Fig. 4k). These data highlight the potential role of local macrophage proliferation in driving the accumulation of SAMs in the fibrotic niche of human chronic liver disease.

### Fibrogenic phenotype of scar-associated macrophages

To delineate the functional profile of SAMs we generated self-organising maps using the *SCRAT* package, visualising co-ordinately expressed gene groups across the MP subpopulations. This created a landscape of 3600 metagenes on a 60×60 grid and highlighted 44 metagene signatures overexpressed in the MP lineage (Fig. 3a). Mapping the nine MP cell clusters to this landscape (Extended Data Fig. 5a) identified six optimally-differentiating metagene signatures, denoted as A-F (Fig. 3a, Supplementary Table 8). Signatures A and B defined the scar-associated macrophages, and were enriched for ontology terms relevant to tissue fibrosis and associated processes such as angiogenesis, in addition to known macrophage-functions such as phagocytosis and antigen processing (Fig. 3b). These SAM-defining signatures included genes such as *TREM2*, *IL1B*, *SPP1*, *LGALS3*, *CXCR4*, *CCR2*, and *TNFSF12*; a number of which are known to regulate the function of scar-producing myofibroblasts in fibrotic liver diseases^19–24^. The remaining MP subpopulations were defined by signature C (Kupffer cells), signatures D, E (tissue monocytes) and signature F (cDC1); ontology terms matched known functions for the associated cell type (Fig. 3b, Extended Data Fig. 5b, Supplementary Table 8). In particular, the Kupffer cell clusters showed significant enrichment for ontology terms involving endocytosis, lipid and iron homeostasis, known functions of KCs in mice^25^. Importantly, macrophage populations did not conform to either an M1 or M2 phenotype, again highlighting the limitation of this classification.

**Figure 5:**
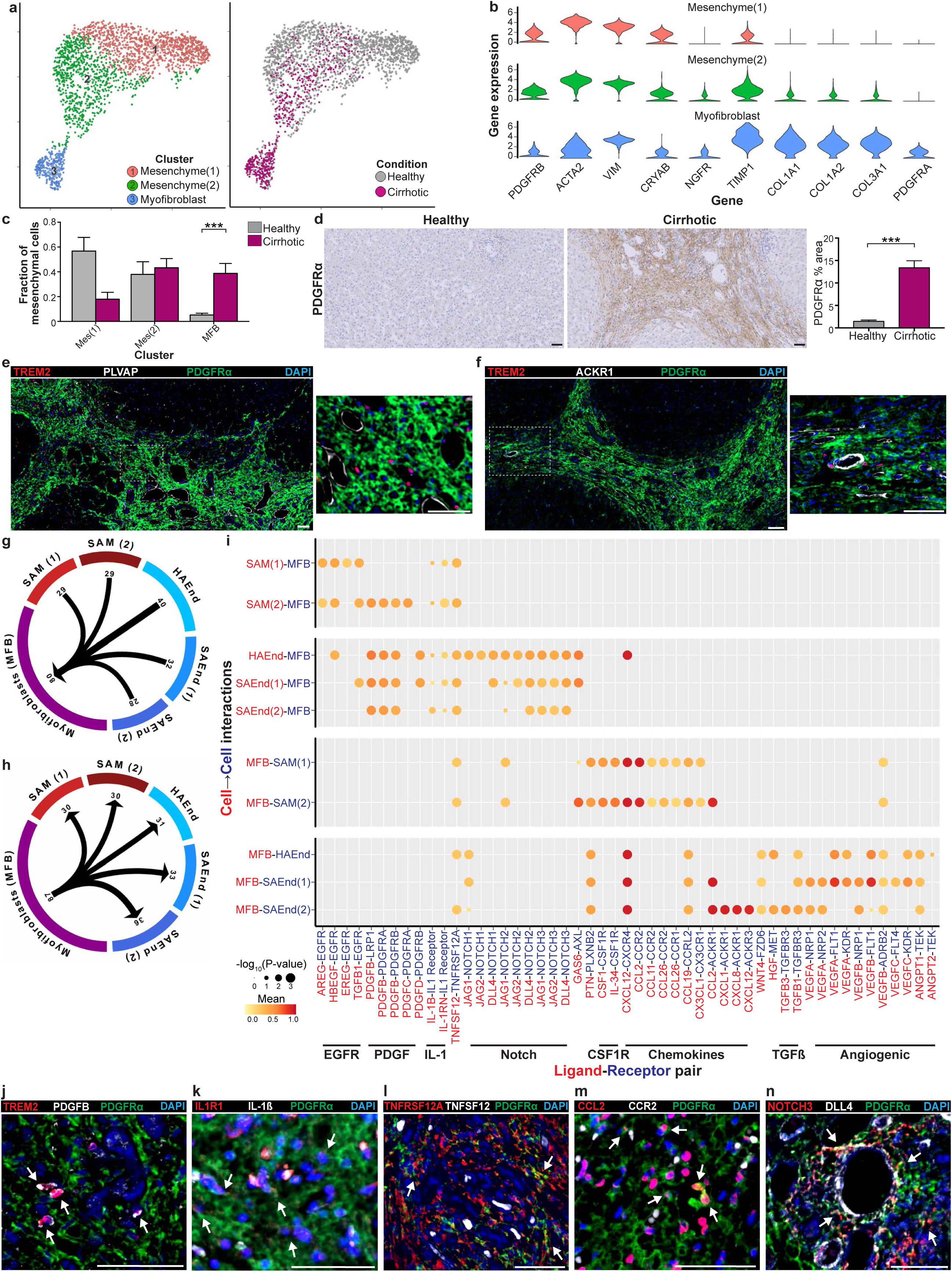
Multi-lineage interactions in the fibrotic niche. **a**, t-SNE visualisation: clustering 2,210 mesenchymal cells, annotating injury condition. **b**, Violin plots: mesenchymal and myofibroblast cluster markers. **c**, Fractions of mesenchymal subpopulations in healthy (n=4) versus cirrhotic (n=3) livers, Wald. **d**, Digital morphometric pixel quantification: PDGFRα staining, healthy (n=11) versus cirrhotic (n=11) liver, Mann-Whitney. **e**, Representative immunofluorescence micrograph, fibrotic niche in cirrhotic liver: TREM2 (red), PLVAP (white), PDGFRα (green), DAPI (blue). **f**, Representative immunofluorescence micrograph, fibrotic niche in cirrhotic liver: TREM2 (red), ACKR1 (white), PDGFRα (green), DAPI (blue). **g**, Circle plot: potential interaction magnitude from ligands expressed by scar-associated macrophages and endothelial cells to receptors expressed on myofibroblasts. **h**, Circle plot: potential interaction magnitude from ligands expressed by myofibroblasts to receptors expressed on scar-associated macrophages and endothelial cells. **i**, Dotplot: selected ligand-receptor interactions between myofibroblasts and scar-associated macrophages and endothelial cells in fibrotic niche. x-axis, ligand (red) and cognate receptor (blue); y-axis, ligand-expressing cell population (red) and receptor-expressing cell population (blue). Circle size indicates p-value, colour (red, high; yellow, low) indicates means of average ligand and receptor expression levels in interacting subpopulations. **j to n**, Representative immunofluorescence micrographs, fibrotic niche in cirrhotic liver. **j**, TREM2 (red), PDGFB (white), PDGFRα (green), DAPI (blue), arrows TREM2^+^PDGFB^+^ cells. **k**, IL1R1 (red), IL-1*β* (white), PDGFRα (green), DAPI (blue), arrows IL1R1^+^PDGFRα^+^ cells. **l**, TNFRSF12A (red), TNFSF12 (white), PDGFRα (green), DAPI (blue), arrows TNFRSF12A^+^PDGFRα^+^ cells. **m**, CCL2 (red), CCR2 (white), PDGFRα (green), DAPI (blue), arrows CCL2^+^PDGFRα^+^ cells. **n**, NOTCH3 (red), DLL4 (white), PDGFRα (green), DAPI (blue), arrows NOTCH3^+^PDGFRα^+^ cells. Scale bars, 50μm. Error bars, s.e.m.; *** p-value < 0.001.

In mice, there are two main origins of hepatic macrophages, either embryologically-derived or monocyte-derived^26^. Under homeostatic conditions, tissue-resident KCs predominate and are embryologically-derived self-renewing cells^27–31^. However, following liver injury, macrophages derived from the recruitment and differentiation of circulating monocytes accumulate in the liver and regulate hepatic fibrosis^14, 32^. The ontogeny of human hepatic macrophage subpopulations has never previously been investigated. Scar-associated TREM2^+^CD9^+^ macrophages demonstrated a monocyte-like morphology (Fig. 2g, Extended Data Fig. 4g-i) and a distinct topographical distribution from KCs (Fig. 2j), suggesting they may represent monocyte-derived cells. To computationally assess the origin of these scar-associated macrophages, we performed *in silico* trajectory analysis on a combined dataset of peripheral blood monocytes and liver-resident MPs. We visualised the transcriptional profile of these cells using a diffusion map, mapped them along a pseudotemporal trajectory (using the *monocle* R package) and interrogated their directionality via spliced and unspliced mRNA ratios (RNA velocity^33, 34^) (Fig. 3c). These analyses suggested a branching differentiation trajectory from peripheral blood monocytes into either scar-associated macrophages or cDCs (Fig. 3c). Additionally, applying RNA velocity indicated a lack of differentiation from KCs to scar-associated macrophages, and no progression from scar-associated macrophages to KCs (Fig. 3c).

To further investigate the pseudotemporal relationship between SAMs and KCs, we visualised the combined blood monocyte and liver-resident MP dataset using a UMAP, and performed additional RNA velocity analyses^33, 34^ (Fig. 3d). Evaluation of spliced and unspliced mRNAs showed expected downregulation (negative velocity) of monocyte gene *MNDA* in SAMs, expected upregulation (positive velocity) of SAM marker gene *CD9* in tissue monocytes, and a lack of KC gene *TIMD4* velocity in SAMs (Fig. 3e). This infers an absence of pseudotemporal dynamics between KCs and SAMs. Furthermore, assessment of the probabilities of cells in this dataset transitioning into SAMs, indicated a higher likelihood of tissue monocytes than KCs differentiating into SAMs (Fig. 3f). Overall, these data suggest that scar-associated macrophages in human fibrotic liver are monocyte-derived, and imply that SAMs represent a terminally-differentiated cell state within the fibrotic niche.

To further characterise the phenotype of scar-associated macrophages, we identified differentially expressed genes along the branching monocyte differentiation trajectories (Fig. 3g). We defined three gene co-expression modules by hierarchical clustering, with module 1 representing genes that are upregulated during blood monocyte-to-SAM differentiation (Fig. 3g). Module 1 was over-expressed in scar-associated macrophages, and contained multiple fibrogenic genes including *SPP1*, *LGALS3*, *CCL2*, *CXCL8*, *PDGFB* and *VEGFA*^19–22,35–37^ (Fig. 3h). Analogous to signatures A and B (Fig. 3b), module 1 displayed ontology terms consistent with promoting tissue fibrosis and angiogenesis, including the regulation of other relevant cell types such as fibroblasts and endothelial cells (Fig. 3h). We confirmed that SAMs show enhanced protein secretion of several of the fibrogenic mediators identified by transcriptional analysis (Extended Data Fig 5c). Co-expression module 2 contained genes that were downregulated during monocyte-to-SAM differentiation, confirming a loss of characteristic monocyte genes (Extended Data Fig. 5d). Module 3 encompassed a distinct group of genes that were upregulated during monocyte-to-cDC differentiation (Extended Data Fig. 5e). Full lists of genes and ontology terms for all three modules are available (Supplementary Table 9). These data highlight that SAMs acquire a specific fibrogenic phenotype during differentiation from circulating monocytes.

To identify potential transcriptional regulators of SAMs we used the *SCENIC* package to define sets of genes co-expressed with known transcription factors, termed regulons. We assessed the cell activity score for differentially-expressed regulons along the tissue monocyte-macrophage pseudotemporal trajectory and in KCs, allowing visualisation of regulon activity across liver-resident macrophage subpopulations (Extended Data Fig. 5f, g, Supplementary Table 10). This identified regulons and corresponding transcription factors associated with distinct macrophage phenotypes, highlighting *NR1H3* and *SPIC* activity in human KCs (Extended Data Fig. 5f, g), which are known to regulate Kupffer cell function in mice^38, 39^. Scar-associated macrophages are enriched for regulons containing the transcription factors *HES1* and *EGR2* (Extended Data Fig. 5f, g), both of which have been associated with modulation of macrophage phenotype and tissue fibrosis^40–43^.

In summary, multimodal computational analysis suggests that TREM2^+^CD9^+^ scar-associated macrophages derive from the recruitment and differentiation of circulating monocytes and display a fibrogenic phenotype.

### Distinct endothelial subpopulations inhabit the fibrotic niche

In rodent models, hepatic endothelial cells are known to regulate both fibrogenesis^44, 45^ and macrophage recruitment to the fibrotic niche^36^. Unsupervised clustering of human liver endothelial cells identified seven subpopulations (Fig. 4a). Clusters Endo(6) and Endo(7) significantly expanded in cirrhotic compared to healthy livers, whilst Endo(1) contracted (Fig. 4a, b). Classical endothelial cell markers did not discriminate between the seven clusters, although Endo(1) was distinct in lacking *CD34* expression (Extended Data Fig. 6a). In order to fully annotate endothelial subpopulations (Extended Data Fig. 6f), we identified differentially expressed markers (Fig. 4c, Supplementary Table 11), determined functional expression profiles (Extended Data Fig. 6c-e, Supplementary Table 12), performed transcription factor regulon analysis (Fig. 4g, Supplementary Table 13) and assessed spatial distribution via multiplex immunofluorescence staining (Fig. 4d, e).

**Figure 6:**
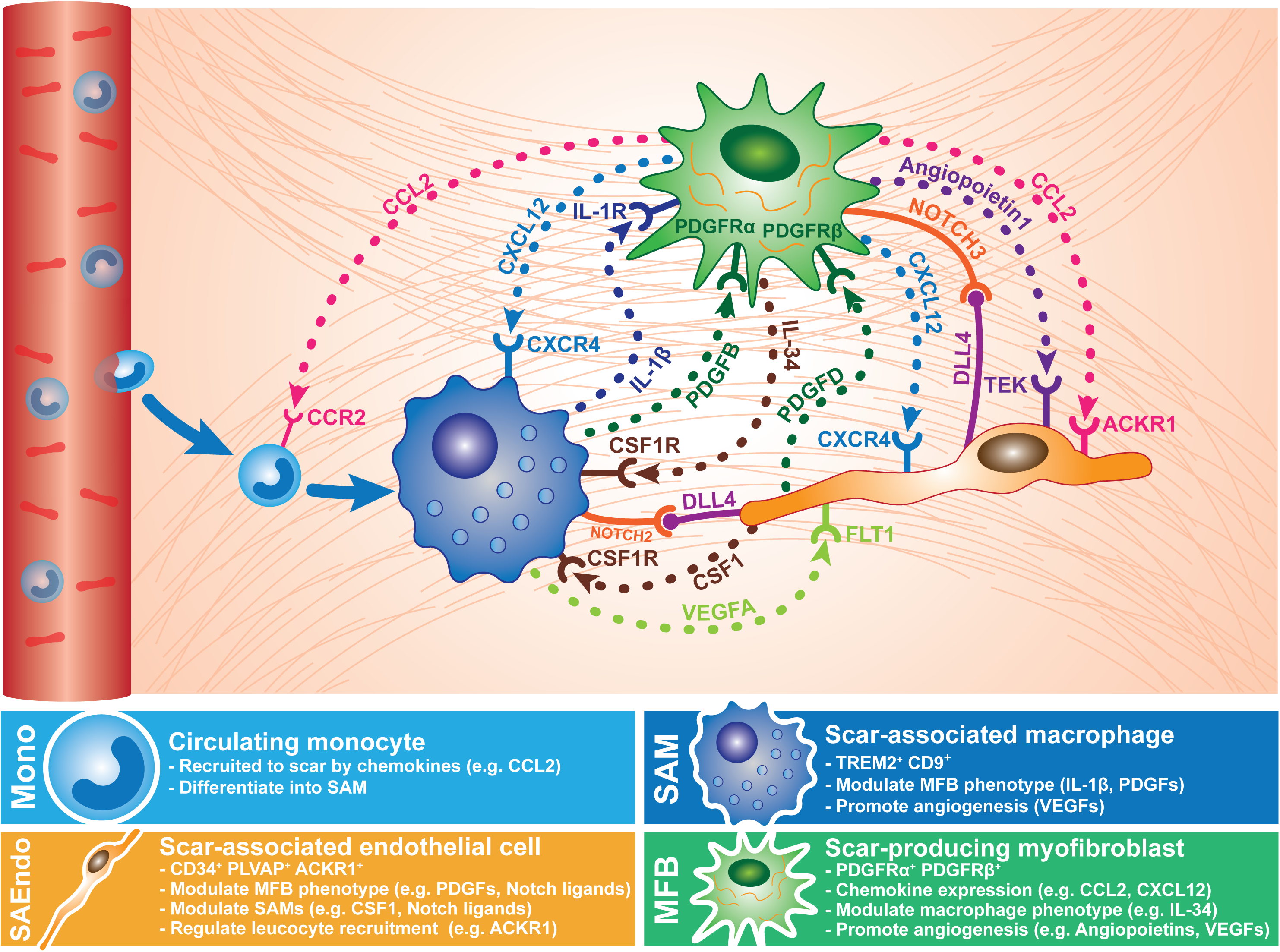
Schematic diagram of cellular interactions in the fibrotic niche of cirrhotic human liver.

Disease-specific endothelial cells Endo(6) and Endo(7), CD34^+^PLVAP^+^ and ACKR1^+^CD34^+^PLVAP^+^ respectively (Fig. 4c, Extended Data Fig. 6b), were spatially restricted to the fibrotic niche in cirrhotic livers (Fig. 4d, e), allowing their annotation as scar-associated endothelia SAEndo(1) and SAEndo(2) respectively. Immunohistochemical quantification of ACKR1 and PLVAP confirmed expansion of these populations in cirrhotic livers, with clear localization within fibrotic septae (Fig. 4f). Scar-associated endothelial cells displayed enhanced expression of the *ELK3* regulon (Fig. 4g), a transcription factor known to modulate angiogenesis^46^. Metagene signature analysis found that Endo(6) (SAEndo(1)) cells expressed fibrogenic genes including *PDGFD*, *PDGFB*, *LOX*, *LOXL2* and several basement membrane components^47–49;^ associated significant ontology terms included extracellular matrix organization and wound healing (signature A; Extended Data Fig. 6c-e). Endo(7) (SAEndo(2)) cells displayed an immunomodulatory phenotype (signature B; Extended Data Fig. 6c-e). Furthermore, the most discriminatory marker for this cluster, ACKR1, is restricted to venules in mice^50^ and has a role in regulating leucocyte recruitment via transcytosis of chemokines from the abluminal to luminal side of blood vessels^51^.

Using *CLEC4M* as a discriminatory marker of cluster Endo(1) (Fig. 4c, Extended Data Fig. 6b), immunofluorescence confirmed these CLEC4M^+^CD34^-^ cells as liver sinusoidal endothelial cells (LSECs), restricted to perisinusoidal cells within the liver parenchyma (Fig. 4d). Cluster Endo(1) demonstrated known features of LSECs, including *GATA4* transcription factor regulon expression^52^ (Fig. 4g), and a metagene signature enriched for ontology terms including endocytosis and immune response^53^ (signature D, Extended Data Fig. 6c-e). There was a reduction in CLEC4M staining in cirrhotic livers with an absence in fibrotic septae (Fig. 4d, f), indicating that LSECs do not inhabit the fibrotic niche in chronic liver disease. This was further supported by trajectory analysis, suggesting a lack of clear pseudotemporal dynamics between the LSECs and clusters SAEndo(1) and SAEndo(2) (Extended Data Fig. 6g).

We annotated cluster Endo(2) (PDPN^+^CD34^+^PLVAP^-^) as lymphatic endothelial cells based on marker gene expression, relevant ontology terms (signature E; Extended Data Fig. 6c-e) and *FOXC2* and *HOXD8* regulon activity (Fig. 4g)^54, 55^. Lymphatics populated the portal region of healthy livers (Fig. 4e). Hierarchical clustering of the endothelial subpopulations demonstrated that clusters Endo(3) and Endo(4) were closely related to LSECs (dendrogram not shown), co-expressing markers including *CLEC4G* (Extended Data Fig. 6b). Endo(4), defined as RSPO3^+^CD34^+^PLVAP^+^ (Fig. 4c, Extended Data Fig. 6b), expressed a metagene signature overlapping with LSECs (signature D, Extended Data Fig. 6c-e), and were identified as central vein endothelial cells (Fig. 4e). This mirrors murine liver zonation data indicating RSPO3 as a marker of pericentral endothelial cells^56^. Similar to LSECs, central vein endothelial cells did not inhabit the fibrotic niche in cirrhosis (Fig. 4e).

Cluster Endo(5), AIF1L^+^CD34^+^PLVAP^+^ cells, were mapped to periportal thick-walled vessels, consistent with hepatic arterial endothelial cells (Fig. 4e). Of note, these cells were also topographically associated with fibrotic septae in cirrhotic livers (Fig. 4e). The arterial identity of this cluster was further indicated by *SOX17* regulon expression^57^ (Fig. 4g), and it displayed a metagene signature enriched for Notch pathway ligands *JAG1*, *JAG2* and *DLL4*; ontology terms included animal organ development, angiogenesis and Notch signalling (signature C; Extended Data Fig. 6c-e) in keeping with the known requirement of the Notch pathway in the development and maintenance of hepatic vasculature^58^. Endo(5) was annotated as HAEndo for subsequent analysis of cellular interactions within the fibrotic niche.

### Resolving the multi-lineage interactome in the fibrotic niche

To investigate how the newly-defined scar-associated macrophage and endothelial cell subpopulations regulate fibrosis, we assessed interactions between these cells and the myofibroblasts, the key scar-producing cells within the fibrotic niche^59, 60^. Myofibroblasts were identified as a distinct subpopulation following clustering of the mesenchymal lineage (Fig. 5a), expressing fibrogenic genes including *COL1A1*, *COL1A2*, *COL3A1* and *TIMP1* in addition to specific markers such as *PDGFRA* (Fig. 5b, Extended Data Fig. 7a, Supplementary Table 14). This myofibroblast cluster was significantly expanded in cirrhotic livers (Fig. 5c), confirmed histologically using PDGFRα immunohistochemistry (Fig. 5d). Multiplex immunofluorescent staining demonstrated the close topographical association of these PDGFRα^+^ myofibroblasts with TREM2^+^ SAMs and both PLVAP^+^ and ACKR^+^ scar-associated endothelial cells within the fibrotic niche (Fig. 5e, f).

To interrogate potential ligand-receptor interactions between these scar-associated macrophage, endothelial and mesenchymal cells, we used CellPhoneDB, a new repository of curated ligand-receptor interactions integrated with a statistical framework. We calculated statistically significant ligand-receptor pairs, based on expression of receptors by one lineage and ligands by another, using empirical shuffling^61^. We focussed further analysis on differentially expressed interactions between pairs of lineages spatially located in the fibrotic niche.

Numerous statistically significant potential interactions were detected between ligands and cognate receptors expressed by scar-associated macrophages, scar-associated endothelial cells and myofibroblasts within the fibrotic niche (Fig. 5g, h, Extended Data Fig. 7b). Selected ligand-receptor pairings between lineages are summarised (Fig. 5i, Extended Data Fig. 7c) and a full list provided (Supplementary Table 15).

The fibrogenic expression profile of SAMs (Fig. 3b, h, Extended Data Fig. 5c), predicts that scar-associated macrophages modulate myofibroblast phenotype within the fibrotic niche. Ligand-receptor analysis provided a molecular framework for these interactions: scar-associated macrophages expressed ligands for Platelet-derived growth factor receptors (PDGFRs), IL-1 receptor, Epidermal growth factor receptor (*EGFR*), and *TNFRSF12A* on myofibroblasts (Fig. 5i), all known to regulate myofibroblast activation, proliferation, survival and promote liver fibrosis in rodent models^19, 24, 47, 62, 63^. Immunofluorescence staining confirmed the close topographical association of PDGFB-PDGFRα, IL1β-IL1R1 and TNFSF12-TNFRSF12A within the fibrotic niche (Fig. 5j, k, l).

Myofibroblasts display a potent immunoregulatory profile, producing cytokines and chemokines such as *CCL2,* which regulates monocyte-macrophage recruitment and phenotype (Fig. 5i, m). We detected highly significant interactions between *CXCL12* produced by myofibroblasts and *CXCR4* expressed by both scar-associated macrophages and endothelial cells (Fig. 5i), confirmed at protein level within the fibrotic septae (Extended Data Fig. 7d). CXCR4 signaling in both macrophages and endothelia have been associated with promoting tissue fibrosis in mice^44, 64^. IL-34, CSF-1 and CX3CL1 interactions with relevant macrophage receptors are known to modulate macrophage function and survival^65, 66^; IL-34 and CSF-1 are potent macrophage mitogens^65^, potentially explaining the increased proliferation observed in SAMs (Extended Data Fig. 4j, k).

Intrahepatic angiogenesis is associated with both degree of liver fibrosis and portal hypertension, a major clinical consequence of liver cirrhosis^67^. Our ligand-receptor analysis confirmed pro-angiogenic interactions within the fibrotic niche, with both scar-associated macrophages and myofibroblasts producing VEGFs, signalling via receptors on scar-associated endothelia (Fig. 5i, Extended Data Fig. 7c). VEGFA expression by SAMs was confirmed histologically (Extended Data Fig. 7e). Myofibroblasts also express angiopoietins (Fig. 5i, Extended Data Fig. 7f), which modulate angiogenic responses in endothelial cells and promote liver fibrosis^68^. Furthermore, SAMs expressed chemokines such as CCL2 and CXCL8 (Extended Data Fig. 5c), interacting with ACKR1 on scar-associated endothelial cells (Extended Data Fig. 7c) and indicative of an additional immunomodulatory role for SAMs.

Scar-associated endothelial cells regulate neighbouring myofibroblasts and macrophages by expressing PDGFR ligands, chemokines, *CSF1* and *CD200* (Fig. 5i, Extended Data Fig. 7c, g). Additionally, endothelial cell interactions showed a pronounced enrichment for Notch signalling, with non-canonical Notch ligands *DLL4*, *JAG1* and *JAG2* interacting with notch receptors *NOTCH2* and *NOTCH3* on myofibroblasts and *NOTCH2* on macrophages (Fig. 5i, Extended Data Fig. 7c). We confirmed the close apposition of DLL4^+^ cells, NOTCH3^+^ myofibroblasts and NOTCH2^+^ SAMs within the fibrotic niche (Fig. 5n, Extended Data Fig. 7h). Notch signalling regulates myofibroblast phenotype and tissue fibrosis^43^, whilst arterial Notch ligand expression regulates monocyte-derived macrophage differentiation and macrophage function in tissue repair^69^. In keeping with active Notch signalling in SAMs, we demonstrate upregulation of the transcription factor regulon *HES1*, a key downstream target of Notch (Fig. 3h, Extended Data Fig. 5f, g), during differentiation from monocytes.

In summary, our unbiased dissection of the key ligand-receptor interactions between novel scar-associated macrophages, endothelial subpopulations and collagen-producing myofibroblasts in the fibrotic niche reveals several major pathways which promote hepatic fibrosis (Fig. 6). Therapeutic targeting of these intra-scar pathways, represents a rational approach for the discovery of novel anti-fibrotic treatments for patients with chronic liver disease.

## Discussion

The fibrotic niche has not previously been defined in human liver. Here, using scRNA-seq and spatial mapping, we resolve the fibrotic niche of human liver cirrhosis, identifying novel pathogenic subpopulations of TREM2^+^CD9^+^ fibrogenic macrophages, ACKR1^+^ and PLVAP^+^ endothelial cells and PDGFRα^+^ collagen-producing myofibroblasts. We dissect a complex, pro-fibrotic interactome between multiple novel scar-associated cells, and identify highly relevant intra-scar pathways that are potentially druggable. This multi-lineage single cell dataset of human liver cirrhosis should serve as a useful resource for the scientific community, and is freely available for interactive browsing at http://www.livercellatlas.mvm.ed.ac.uk.

Despite significant progress in our understanding of the molecular pathways driving liver fibrosis in rodent models, a lack of corollary studies in diseased human liver tissue has hindered translation into effective therapies, with currently no FDA or EMA-approved anti-fibrotic treatments available. Our multi-lineage ligand-receptor analysis demonstrates the complexity of interactions within the fibrotic niche, highlighting why current approaches to treat human liver fibrosis have proven so intractable, and provides a conceptual framework for more rational studies of anti-fibrotic therapies in both pre-clinical animal models and translational systems such as human liver organoid cultures^5, 70, 71^. Further, this unbiased multi-lineage approach should inform the design of combination therapies which will very likely be necessary to achieve effective anti-fibrotic potency^5, 6^.

Macrophages and endothelial cells are known to regulate liver fibrosis in rodent models^13, 19, 23, 44, 45^. However, little is known regarding the heterogeneity and precise molecular definitions of these cell types in human liver disease. Our data demonstrates both the accumulation of discrete monocyte-derived macrophage and endothelial cell populations in the fibrotic niche of cirrhotic livers, but also the persistence of spatially distinct, non-scar associated resident Kupffer cells and liver sinusoidal endothelial cells. This single-cell approach has important implications for therapy development; facilitating specific targeting of pathogenic cells without perturbing homeostatic function.

In this era of precision medicine, where molecular profiling guides the development of highly targeted therapies, we used scRNA-seq to resolve the key non-parenchymal cell subclasses inhabiting the fibrotic niche of human liver cirrhosis. Application of our novel scar-associated cell markers could potentially inform molecular pathology-based patient stratification, which is fundamental to the prosecution of successful anti-fibrotic clinical trials. Our work illustrates the power of single-cell transcriptomics to decode the cellular and molecular basis of human organ fibrosis, providing a conceptual framework for the discovery of relevant and rational therapeutic targets to treat patients with a broad range of fibrotic diseases.

## Methods

### Study subjects

Local approval for procuring human liver tissue and blood samples for scRNA-seq, flow cytometry and histological analysis was obtained from the NRS BioResource and Tissue Governance Unit (Study Number SR574), following review at the East of Scotland Research Ethics Service (Reference 15/ES/0094). All subjects provided written informed consent. Healthy background non-lesional liver tissue was obtained intraoperatively from patients undergoing surgical liver resection for solitary colorectal metastasis at the Hepatobiliary and Pancreatic Unit, Department of Clinical Surgery, Royal Infirmary of Edinburgh. Patients with a known history of chronic liver disease, abnormal liver function tests or those who had received systemic chemotherapy within the last four months were excluded from this cohort. Cirrhotic liver tissue was obtained intraoperatively from patients undergoing orthotopic liver transplantation at the Scottish Liver Transplant Unit, Royal Infirmary of Edinburgh. Blood from patients with a confirmed diagnosis of liver cirrhosis were obtained from patients attending the Scottish Liver Transplant Unit, Royal Infirmary of Edinburgh. Patients with liver cirrhosis due to viral hepatitis were excluded from the study. Patient demographics are summarised in Extended Data Fig. 1a. Sorting of macrophage subpopulations from cirrhotic livers for analysis of secreted mediators was performed at the University of Birmingham, UK. Local ethical approval was obtained (Reference 06/Q2708/11) and all patients provided written, informed consent. Liver tissue was acquired from explanted diseased livers from patients undergoing orthotopic liver transplantation at the Queen Elizabeth Hospital, Birmingham.

### Tissue processing

For liver scRNA-seq and flow cytometry analysis, a wedge biopsy of non-ischaemic fresh liver tissue (2-3 grams) was obtained by the operating surgeon, prior to interruption of the hepatic vascular inflow. This was immediately placed in HBSS (Gibco) on ice. The tissue was then transported directly to the laboratory and dissociation routinely commenced within 20 minutes of the liver biopsy. To enable paired histological assessment, a segment of each liver specimen was also fixed in 4% neutral-buffered formalin for 24 hours followed by paraffin-embedding. Additional liver samples, obtained via the same method, were fixed in an identical manner and used for further histological analysis. For cell sorting to assess secreted mediator production, explanted diseased liver tissue (40 grams) was used from patients undergoing orthotopic liver transplantation.

### Immunohistochemistry, immunofluorescence and smFISH

Formalin-fixed paraffin-embedded human liver tissue was cut into 4 μm sections, dewaxed, rehydrated, then incubated in 4% neutral-buffered formalin for 20 minutes. Following heat-mediated antigen retrieval in pH6 sodium citrate (microwave; 15 minutes), slides were washed in PBS and incubated in 4% hydrogen peroxide for 10 minutes. Slides were then washed in PBS, blocked using protein block (GeneTex, GTX30963) for 1 hour at room temperature before incubation with primary antibodies for 1 hour at room temperature. A full list of primary antibodies and conditions are shown in Supplementary Table 16. Slides were washed in PBST (PBS plus 0.1% Tween20; Sigma-Aldrich, P1379) then incubated with ImmPress HRP Polymer Detection Reagents (depending on species of primary; rabbit, MP-7401; mouse, MP-6402-15; goat, MP-7405; all Vector Laboratories) for 30 minutes at room temperature. Slides were washed in PBS followed by detection. For DAB staining, sections were incubated with DAB (DAKO, K3468) for 5 minutes and washed in PBS before a haematoxylin (Vector Laboratories, H3404) counterstain. For multiplex immunofluorescence staining, following the incubation with ImmPress and PBS wash, initial staining was detected using either Cy3, Cy5, or Fluorescein tyramide (Perkin-Elmer, NEL741B001KT) at a 1:1000 dilution. Slides were then washed in PBST followed by further heat treatment with pH6 sodium citrate (15 minutes), washes in PBS, protein block, incubation with the second primary antibody (incubated overnight at 4°C), ImmPress Polymer and tyramide as before. This sequence was repeated for the third primary antibody (incubated at room temperature for 1 hour) and a DAPI-containing mountant was then applied (ThermoFisher Scientific, P36931).

For AMEC Staining (only CLEC4M immunohistochemistry), all washes were carried out with TBST (dH2O, 2oomM Tris, 1.5M NaCl, 1% Tween20 (all Sigma-Aldrich) pH7.5) and peroxidase blocking was carried out for 30mins in 0.6% hydrogen peroxide in Methanol. Sections were incubated with AMEC (Vector Laboratories, SK-4285) for 20 minutes and washed in TBST (dH2O, 200mM Tris, 1.5M NaCl, 1% Tween20 (all Sigma-Aldrich)) before a haematoxylin (Vector Laboratories, SK-4285) counterstain.

For combined single-molecule fluorescent in situ hybridization (smFISH) and immunofluorescence, detection of TREM2 was performed using the RNAscope^®^ 2.5 LS Reagent Kit - BrownAssay (Advanced Cell Diagnostics (ACD)) in accordance with the manufacturer’s instructions. Briefly, 5 μm tissue sections were dewaxed, incubated with endogenous enzyme block, boiled in pretreatment buffer and treated with protease, followed by target probe hybridization using the RNAscope^®^ LS 2.5 Hs-TREM2 (420498, ACD) probe. Target RNA was then detected with Cy3 tyramide (Perkin-Elmer, NEL744B001KT) at 1:1000 dilution. The sections were then processed through a pH6 sodium citrate heat-mediated antigen retrieval, hydrogen peroxidase treatment and protein block (all as for multiplex immunofluorescence staining as above). MNDA antibody was applied overnight at 4°C, completed using a secondary ImmPress HRP Anti-Rabbit Peroxidase IgG (Vector Laboratories, MP7401), visualised using a Flourescein tyramide (Perkin-Elmer, NEL741B001KT) at a 1:1000 dilution and stained with DAPI.

Brightfield and fluorescently-stained sections were imaged using the slide scanner AxioScan.Z1 (Zeiss) at 20X magnification (40X magnification for smFISH). Images were processed and scale bars added using Zen Blue (Zeiss) and Fiji software^72^.

### Cell counting and image analysis

Automated cell counting was performed using QuPath software^73^. Briefly, DAB-stained whole tissue section slide-scanned images (CZI files) were imported into QuPath. Cell counts were carried out using the positive cell detection tool, detecting haematoxylin-stained nuclei and then thresholding for positively-stained DAB cells, generating DAB-positive cell counts/mm^2^ tissue. Identical settings and thresholds were applied to all slides for a given stain and experiment. For cell counts of fibrotic septae vs parenchymal nodules, the QuPath segmentation tool was used to segment the DAB-stained whole tissue section into fibrotic septae or non-fibrotic parenchymal nodule regions using tissue morphological characteristics (Fig. 2j). Positive cell detection was then applied to the fibrotic and non-fibrotic regions in turn, providing cell DAB-positive cell counts/mm^2^ in fibrotic septae and non-fibrotic parenchymal nodules for each tissue section.

Digital morphometric pixel analysis was performed using the Trainable Weka Segmentation (TWS) plugin^74^ in Fiji software^72^. Briefly, each stained whole tissue section slide-scanned image was converted into multiple TIFF files in Zen Blue software (Zeiss). TIFF files were imported into Fiji and TWS plugin trained to produce a classifier which segments images into areas of positive staining, tissue background and white space^74^. The same trained classifier was then applied to all TIFF images from every tissue section for a particular stain, providing a percentage area of positive staining for each tissue section.

### Preparation of single-cell suspensions

For liver scRNA-seq, human liver tissue was minced with scissors and digested in 5mg/ml pronase (Sigma-Aldrich, P5147-5G), 2.93mg/ml collagenase B (Roche, 11088815001) and 1.9mg/ml DNase (Roche, 10104159001) at 37°C for 30 minutes with agitation (200–250 r.p.m.), then strained through a 120μm nybolt mesh along with PEB buffer (PBS, 0.1% BSA, and 2mM EDTA) including DNase (0.02mg/ml). Thereafter all processing was done at 4°C. The cell suspension was centrifuged at 400*g* for 7 minutes, supernatant removed, cell pellet resuspended in PEB buffer and DNase added (0.02mg/ml), followed by additional centrifugation (400g, 7 minutes). Red blood cell lysis was performed (BioLegend, 420301), followed by centrifugation (400*g*, 7 minutes), resuspension in PEB buffer and straining through a 35μm filter. Following another centrifugation at 400*g* for 7 minutes, cells were blocked in 10% human serum (Sigma-Aldrich, H4522) for 10 minutes at 4°C prior to antibody staining.

For both liver flow cytometry analysis and cell sorting to assess secreted mediator production, single-cell suspensions were prepared as previously described, with minor modifications^75^. In brief, liver tissue was minced and digested in an enzyme cocktail 0.625 mg/ml collagenase D (Roche, 11088882001), 0.85 mg/ml collagenase V (Sigma-Aldrich, C9263-1G), 1 mg/ml dispase (Gibco, Invitrogen, 17105-041), and 30 U/ml DNase (Roche, 10104159001) in RPMI-1640 at 37°C for 45 minutes with agitation (200–250 r.p.m.), before being passed through a 100μm filter. Following red blood cell lysis (BioLegend, 420301), cells were passed through a 35μm filter. Before the addition of antibodies, cells were blocked in 10% human serum (Sigma-Aldrich, H4522) for 10 minutes at 4 °C.

For PBMC scRNA-seq, 4.9ml peripheral venous blood samples were collected in EDTA-coated tubes (Sarstedt, S-Monovette*®* 4.9ml K3E) and placed on ice. Blood samples were transferred into a 50ml Falcon tube. Following red cell lysis (Biolegend, 420301), blood samples were then centrifuged at 500g for 5 minutes and supernatant was removed. Pelleted samples were then resuspended in staining buffer (PBS plus 2% BSA; Sigma-Aldrich) and centrifugation was repeated. Samples were then blocked in 10% human serum (Sigma-Aldrich, H4522) in staining buffer on ice for 30 minutes. Cells were then resuspended in staining buffer and passed through a 35μm filter prior to antibody staining.

### Flow cytometry and cell sorting

Incubation with primary antibodies was performed for 20 minutes at 4°C. All antibodies, conjugates, lot numbers and dilutions used in this study are presented in Supplementary Table 16. Following antibody staining, cells were washed with PEB buffer. For both flow cytometry analysis and cell sorting to assess secreted mediator production, cells were then incubated with streptavidin-BV711 for 20 minutes at 4°C (Biolegend 405241; Dilution 1:200).

For cell sorting (FACS), cell viability staining (DAPI; 1:1000 dilution) was then performed, immediately prior to acquiring the samples. Cell sorting for scRNA-seq was performed on a BD Influx (Becton Dickinson, Basel, Switzerland). Cell sorting to assess secreted mediator production was performed on a BD FACSAria^TM^ Fusion (Becton Dickinson, Basel, Switzerland).

For flow cytometry analysis, cells were then stained with Zombie NIR fixable viability dye (Biolegend, 423105) according to manufacturers’ instructions. Cells were washed in PEB then fixed in IC fixation buffer (Thermo-Fisher, 00-8222-49) for 20 minutes at 4°C. Fixed samples were stored in PEB at 4°C until acquisition. Flow cytometry acquistion was performed on 6-laser Fortessa flow cytometer (Becton Dickinson, Basel, Switzerland) and data analyzed using Flowjo software (Treestar, Ashland, TN).

### Analysis of secreted mediators

To assess secreted mediators produced by macrophage subpopulations from cirrhotic livers, FACS-sorted SAMs (Viable CD45^+^Lin^-^HLA-DR^+^CD14^+^CD16^+^CD163^-^ TREM2^+^CD9^+^), TMo (Viable CD45^+^Lin^-^ HLA-DR^+^CD14^+^CD16^+^CD163^-^) and KCs (Viable CD45^+^Lin^-^ HLA-DR^+^CD14^+^CD16^+^CD163^+^CD9^-^) were cultured in 12-well plates (Corning, 3513) in DMEM (Gibco, 41965039) containing 2% FBS (Gibco, 10500056) at 1×10^6^ cells/ml for 24 hours at 37°C. Control wells contained media alone. Conditioned media was collected, centrifuged at 400g for 10 minutes and stored at −80°C. Detection of CCL2, CD163, Galectin-3, IL-1 beta, CXCL8 and Osteopontin (SPP1) proteins in conditioned media was performed using a custom human luminex assay (R&D systems), according to the manufacturers protocol. Data was acquired using a Bio-Plex*^®^* 200 (Bio-Rad, UK) and is presented a median fluorescence intensity (MFI) for each analyte.

### Droplet-based scRNA-seq

Single cells were processed through the Chromium*™* Single Cell Platform using the Chromium*™* Single Cell 3’ Library and Gel Bead Kit v2 (10X Genomics, PN-120237) and the Chromium*™* Single Cell A Chip Kit (10X Genomics, PN-120236) as per the manufacturer’s protocol. In brief, single cells were sorted into PBS + 0.1% BSA, washed twice and counted using a Bio-Rad TC20. 10,769 cells were added to each lane of the 10X chip. The cells were then partitioned into Gel Beads in Emulsion in the Chromium*™* instrument, where cell lysis and barcoded reverse transcription of RNA occurred, followed by amplification, fragmentation and 5′ adaptor and sample index attachment. Libraries were sequenced on an Illumina HiSeq 4000.

### Pre-processing scRNA-seq data

We aligned to the GRCh38 reference genome, and estimated cell-containing partitions and associated UMIs, using the Cell Ranger v2.1.0 Single-Cell Software Suite from 10X Genomics. Genes expressed in fewer than three cells in a sample were excluded, as were cells that expressed fewer than 300 genes or mitochondrial gene content >30% of the total UMI count. We normalised by dividing the UMI count per gene by the total UMI count in the corresponding cell and log-transforming. Variation in UMI counts between cells was regressed according to a negative binomial model, before the resulting value was scaled and centred by subtracting the mean expression of each gene and dividing by its standard deviation (En), then calculating ln(10^4^*En+1).

### Dimensionality reduction, clustering, and DE analysis

We performed unsupervised clustering and differential gene expression analyses in the *Seurat* R package v2.3.0^76^. In particular we used SNN graph-based clustering, where the SNN graph was constructed using between 2 and 11 principal components as determined by dataset variability shown in principal components analysis (PCA); the resolution parameter to determine the resulting number of clusters was also tuned accordingly. To assess cluster similarity we used the *BuildClusterTree* function from *Seurat*.

In total, we present scRNA-seq data from ten human liver samples (named Healthy 1-5 and Cirrhotic 1-5) and five human blood samples (n=4 cirrhotic named Blood 1-4 and n=1 healthy named PBMC8K; pbmc8k dataset sourced from single-cell gene expression datasets hosted by 10X Genomics). For seven liver samples (Healthy 1-4 and Cirrhotic 1-3) we performed scRNA-seq on both leucocytes (CD45^+^) and other non-parenchymal cells (CD45^-^); for the remaining three livers (Healthy 5, Cirrhotic 4-5) we performed scRNA-seq on leucocytes only (Extended Data Fig. 2e, g).

Initially, we combined all scRNA-seq datasets (liver and blood) and performed clustering analysis with the aim of isolating a population of liver-resident cells, by identifying contaminating circulatory cells within datasets generated from liver digests and removing them from downstream analysis. Specifically, we removed from our liver datasets cells that fell into clusters 1 and 13 of the initial dataset in Extended Data Fig. 1d.

Using further clustering followed by signature analysis, we interrogated this post-processed liver-resident cell dataset for robust cell lineages. These lineages were isolated into individual datasets, and the process was iterated to identify robust lineage subpopulations. At each stage of this process we removed clusters expressing more than one unique lineage signature in more than 25% of their cells from the dataset as probable doublets. Where the cell proliferation signature identified distinct cycling subpopulations, we re-clustered these again to ascertain the identity of their constituent cells.

All heatmaps, t-SNE and UMAP visualisations, violin plots, and dot plots were produced using *Seurat* functions in conjunction with the *ggplot2*, *pheatmap*, and *grid* R packages. t-SNE and UMAP visualisations were constructed using the same number of principal components as the associated clustering, with perplexity ranging from 30 to 300 according to the number of cells in the dataset or lineage. We conducted differential gene expression analysis in *Seurat* using the standard AUC classifier to assess significance. We retained only those genes with a log-fold change of at least 0.25 and expression in at least 25% of cells in the cluster under comparison.

### Defining cell lineage signatures

For each cell we obtained a signature score across a curated list of known marker genes per cell lineage in the liver (Supplementary Table 1). This signature score was defined as the geometric mean of the expression of the associated signature genes in that cell. Lineage signature scores were scaled from 0 to 1 across the dataset, and the score for each cell with signature less than a given threshold (the mean of said signature score across the entire dataset) was set as 0.

### Batch effect and quality control

To investigate agreement between samples we extracted the average expression profile for a given cell lineage in each sample, and calculated the Pearson correlation coefficients between all possible pairwise comparisons of samples per lineage^77^.

### Imputing dropout in T cell and ILC clusters

To impute dropout of low-abundance transcripts in our T cell and ILC clusters so that we might associate them with known subpopulations, we down-sampled to 7,380 cells from 36,900 and applied the *scImpute* R package v0.0.8^78^, using as input both our previous annotation labels and k-means spectral clustering (k=5), but otherwise default parameters.

### Analysing functional phenotypes of scar-associated cells

For further analysis of function we adopted the self-organising maps (SOM) approach as implemented in the *SCRAT* R package v1.0.0^79^. For each lineage of interest we constructed a SOM in *SCRAT* using default input parameters and according to its clusters. We defined the signatures expressed in a cell by applying a threshold criterion (e^thresh^ = 0.95 × e^max^) selecting the highest-expressed metagenes in each cell, and identified for further analysis those metagene signatures defining at least 30% of cells in at least one cluster within the lineage. We smoothed these SOMs using the *disaggregate* function from the *raster* R package for visualisation purposes, and scaled radar plots to maximum proportional expression of the signature. Gene ontology enrichment analysis on the genes in these spots was performed using PANTHER 13.1 (pantherdb.org).

### Inferring injury dynamics and transcriptional regulation

To generate cellular trajectories (pseudotemporal dynamics) we used the *monocle* R package v2.6.1^80^. We ordered cells in a semi-supervised manner based on their *Seurat* clustering, scaled the resulting pseudotime values from 0 to 1, and mapped them onto either the t-SNE or UMAP visualisations generated by *Seurat* or diffusion maps as implemented in the *scater* R package v1.4.0^81^ using the top 500 variable genes as input. We removed mitochondrial and ribosomal genes from the geneset for the purposes of trajectory analysis. Differentially-expressed genes along this trajectory were identified using generalised linear models via the *differentialGeneTest* function in *monocle*.

When determining significance for differential gene expression along the trajectory, we set a q-value threshold of 1e-20. We clustered these genes using hierarchical clustering in *pheatmap*, cutting the tree at k=3 to obtain gene modules with correlated gene expression across pseudotime. Cubic smoothing spline curves were fitted to scaled gene expression along this trajectory using the *smooth.spline* command from the *stats* R package, and gene ontology enrichment analysis again performed using PANTHER 13.1.

We verified the trajectory and its directionality using the *velocyto* R package v0.6.0^33^, estimating cell velocities from their spliced and unspliced mRNA content. We generated annotated spliced and unspliced reads from the 10X BAM files via the *dropEst* pipeline, before calculating gene-relative velocity using kNN pooling with k=25, determining slope gamma with the entire range of cellular expression, and fitting gene offsets using spanning reads. Aggregate velocity fields (using Gaussian smoothing on a regular grid) and transition probabilities per lineage subpopulations were visualised on t-SNE, UMAP, or diffusion map visualisations as generated previously. Gene-specific phase portraits were plotted by calculating spliced and unspliced mRNA levels against steady-state inferred by a linear model; levels of unspliced mRNA above and below this steady-state indicate increasing and decreasing expression of said gene, respectively. Similarly we plotted unspliced count signal residual per gene, based on the estimated gamma fit, with positive and negative residuals indicating expected upregulation and downregulation respectively.

For transcription factor analysis, we obtained a list of all genes identified as acting as transcription factors in humans from AnimalTFDB^82^. To further analyse transcription factor regulons, we adopted the *SCENIC* v0.1.7 workflow in R^83^, using default parameters and the normalised data matrices from *Seurat* as input. For visualisation, we mapped the regulon activity (AUC) scores thus generated to the pseudotemporal trajectories from *monocle* and the clustering subpopulations from *Seurat*.

### Analysing inter-lineage interactions within the fibrotic niche

For comprehensive systematic analysis of inter-lineage interactions within the fibrotic niche we used CellPhoneDB^61^. CellPhoneDB is a manually curated repository of ligands, receptors, and their interactions, integrated with a statistical framework for inferring cell-cell communication networks from single-cell transcriptomic data. In brief, we derived potential ligand-receptor interactions based on expression of a receptor by one lineage subpopulation and a ligand by another; as input to this algorithm we used cells from the fibrotic niche as well as liver sinusoidal endothelial cells and Kupffer cells as control, and we considered only ligands and receptors expressed in greater than 5% of the cells in any given subpopulation. Subpopulation-specific interactions were identified as follows: 1) randomly permuting the cluster labels of all cells 1000 times and determining the mean of the average receptor expression of a subpopulation and the average ligand expression of the interacting subpopulation, thus generating a null distribution for each ligand-receptor pair in each pairwise comparison between subpopulations, 2) calculating the proportion of these means that were “as or more extreme” than the actual mean, thus obtaining a p-value for the likelihood of subpopulation specificity for a given ligand-receptor pair, 3) prioritising interactions that displayed specificity to subpopulations interacting within the fibrotic niche.

### Statistical Analysis

To assess whether our identified subpopulations were significantly overexpressed in injury, we posited the proportion of injured cells in each cluster as a random count variable using a Poisson process, as previously described^77^. We modelled the rate of detection using the total number of cells in the lineage profiled in a given sample as an offset, with the condition of each sample (healthy vs cirrhotic) provided as a covariate factor. The model was fitted using the R command *glm* from the *stats* package. The p-value for the significance of the proportion of injured cells was assessed using a Wald test on the regression coefficient. This methodology was also applied to assess significant changes in proportions of mononuclear phagocytes between healthy and cirrhotic liver tissue by flow cytometry.

Remaining statistical analyses were performed using GraphPad Prism (GraphPad Software, USA). Comparison of changes in histological cell counts or morphometric pixel analysis between healthy and cirrhotic livers were performed using a Mann-Whitney test (unpaired; two-tailed). Comparison of topographical localisation of counted cells (fibrotic septae vs parenchymal nodule) was performed using a Wilcoxon matched-pairs signed rank test (paired; two-tailed). P-values<0.05 were considered statistically significant.

## Data and materials availability

Our expression data will be freely available for user-friendly interactive browsing online http://www.livercellatlas.mvm.ed.ac.uk. CellPhoneDB^61^ is available at www.CellPhoneDB.org, along with lists of membrane proteins, ligands and receptors, and heteromeric complexes. All raw sequencing data is available in the Gene Expression Omnibus (GEO accession GSE136103).

## Code availability

R scripts enabling the main steps of the analysis are available from the corresponding authors on reasonable request.

## Acknowledgements

This work was supported by an MRC Clinician Scientist Fellowship (MR/N008340/1) to P.R., a Wellcome Trust Senior Research Fellowship in Clinical Science (ref. 103749) to N.C.H., an AbbVie Future Therapeutics and Technologies Division grant to N.C.H., a Guts UK– Children’s Liver Disease Foundation grant (ref. R43927) to N.C.H., a Tenovus Scotland grant (ref. E18/05) to R.D. and N.C.H., and British Heart Foundation grants (RM/17/3/33381; RE/18/5/34216) to N.C.H. R.V-T. was funded by EMBO and Human Frontiers long-term fellowships. C.J.W. was funded by a BBSRC New Investigator Award (BB/N018869/1). P.N.N., C.J.W. and N.T.L. are funded by the NIHR Birmingham Biomedical Research Centre at the University Hospitals Birmingham NHS Foundation Trust and the University of Birmingham. This paper presents independent research supported by the NIHR Birmingham Biomedical Research Centre at the University Hospitals Birmingham NHS Foundation Trust and the University of Birmingham. The views expressed are those of the author(s) and not necessarily those of the NHS, the NIHR or the Department of Health and Social Care. J.P.I. was funded by the NIHR Bristol Biomedical Research Centre, University Hospitals Bristol Foundation Trust and the University of Bristol. C.P.P. was funded by the UK Medical Research Council, MC_UU_00007/15. This work was also supported by Wellcome Sanger core funding (WT206194). We thank the patients who donated liver tissue and blood for this study. We thank J. Davidson, C. Ibbotson, J. Black and A. Baird of the Scottish Liver Transplant Unit and the research nurses of the Wellcome Trust Clinical Research Facility for assistance with consenting patients for this study. We thank the liver transplant coordinators and surgeons of the Scottish Liver Transplant Unit and the surgeons and staff of the Hepatobiliary Surgical Unit, Royal Infirmary of Edinburgh for assistance in procuring human liver samples. We thank S. Johnston, W. Ramsay and M. Pattison (QMRI Flow Cytometry and Cell Sorting Facility, University of Edinburgh) for technical assistance with fluorescence activated cell sorting (FACS) and flow cytometry. This publication is part of the Human Cell Atlas (www.humancellatlas.org/publications).

## Author Contributions

P.R. performed experimental design, data generation, data analysis and interpretation, and manuscript preparation; R.D., E.D., K.P.M., B.E.P.H. and N.T.L. assisted with experiments; J.R.P. generated the interactive online browser for our single cell transcriptomic data; M.E. and R.V-T. assisted with CellPhoneDB analyses and critically appraised the manuscript; C.J.W. and P.N.N. provided intellectual contribution; J.R.W-K. performed computational analysis with assistance from R.S.T. and advice from C.P.P., J.M. and S.A.T.; J.W-K. also helped with manuscript preparation and C.P.P., J.M. and S.A.T. critically appraised the manuscript; E.M.H., D.J.M. and S.J.W. procured human liver tissue and critically appraised the manuscript. J.P.I., F.T. and J.W.P. provided intellectual contribution and critically appraised the manuscript; N.C.H. conceived the study, designed experiments, interpreted data and prepared the manuscript.

## Author Information

The authors declare no competing financial interests. Address correspondence or requests for materials to Prakash Ramachandran or Neil Henderson.

---

**Extended Data Figure 1:**
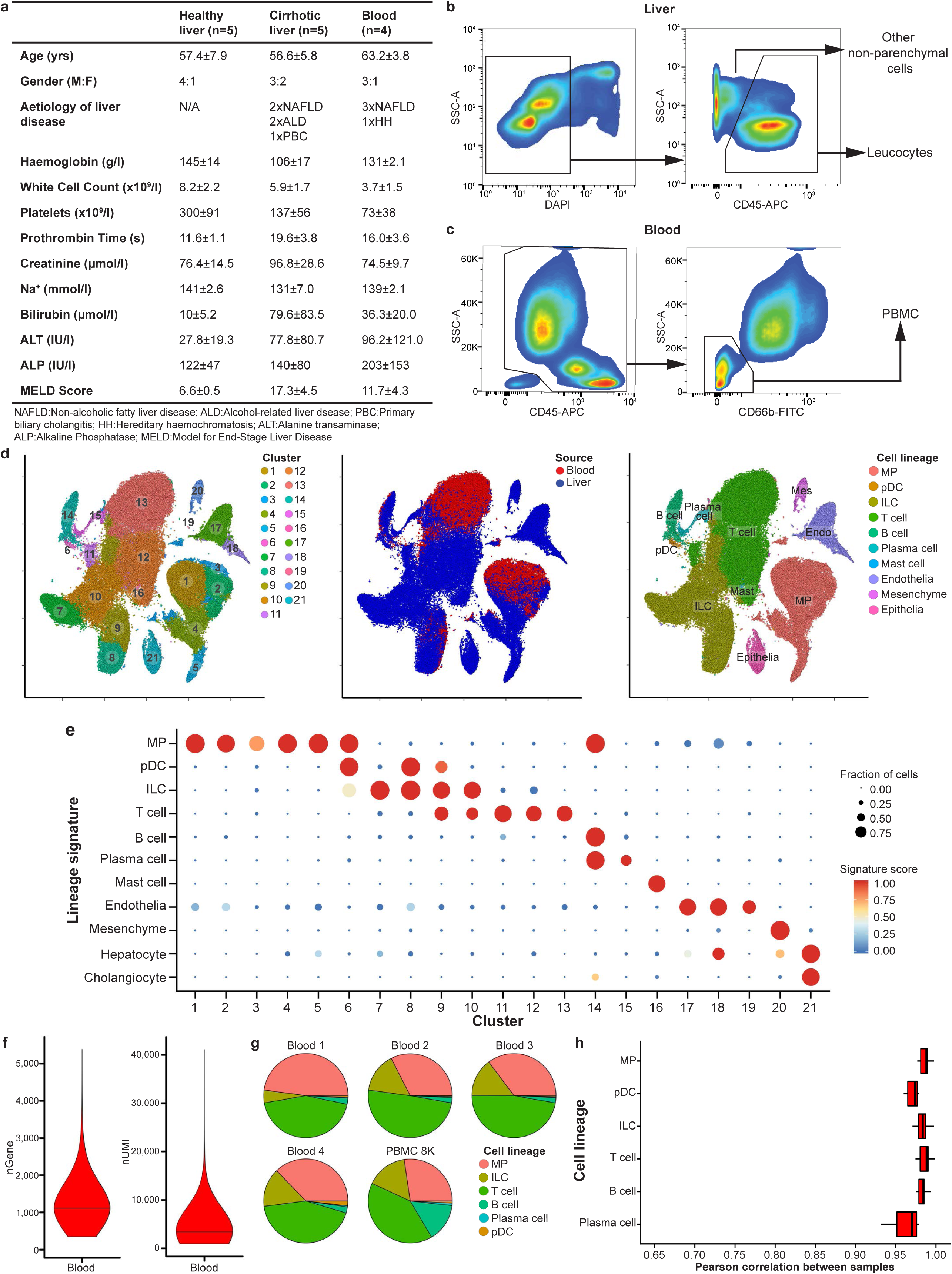
Strategy for isolation of human liver non-parenchymal cells. a, Patient demographics and clinical information. b, Representative flow cytometry plots: gating strategy for isolating leucocytes (CD45^+^) and other non-parenchymal cells (CD45^-^) from healthy and cirrhotic liver. c, Representative flow cytometry plots: gating strategy for isolating peripheral blood mononuclear cells (PBMC). d, t-SNE visualisation: clustering 103,568 cells (n=5 healthy human livers, n=5 cirrhotic human livers, n=1 healthy PBMC, n=4 cirrhotic PBMC), annotating source (PBMC versus liver) and cell lineage inferred from known marker gene signatures. Endo, endothelial cell; ILC, innate lymphoid cell; Mast, mast cell; Mes, mesenchymal cell; MP, mononuclear phagocyte; pDC, plasmacytoid dendritic cell. e, Dotplot: annotating PBMC and liver dataset clusters by lineage signatures. Circle size indicates cell fraction expressing signature greater than mean; colour indicates mean signature expression (red, high; blue, low). f, Violin plots: number of unique genes (nGene) and number of total Unique Molecular Identifiers (nUMI) expressed in PBMC. g, Pie charts: proportion of cell lineage per PBMC sample. h, Box and whisker plot: agreement in expression profiles across PBMC samples. Pearson correlation coefficients between average expression profiles for cell in each lineage, across all pairs of samples. Black bar, median value; box edges, 25^th^ and 75^th^ percentiles; whiskers, full range.

**Extended Data Figure 2:**
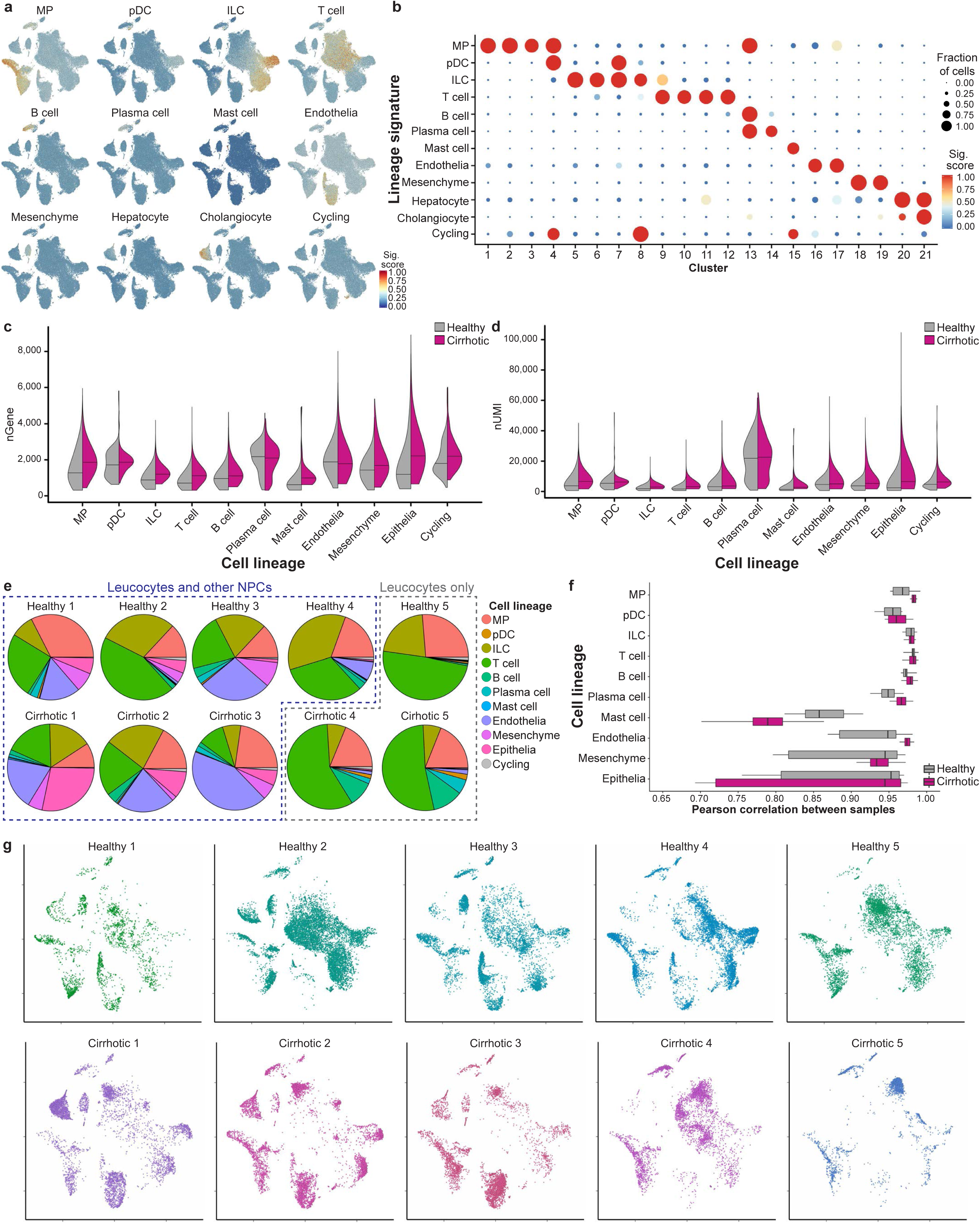
Quality control and annotation of human liver-resident cells. **a**, t-SNE visualisation: lineage signature expression across liver-resident cell dataset (red, high; blue, low). **b**, Dotplot: annotating liver-resident cell clusters by lineage signature. Circle size indicates cell fraction expressing signature greater than mean; colour indicates mean signature expression (red, high; blue, low). **c**, Violin plot: number of unique genes (nGene) expressed across liver-resident cell lineages in healthy versus cirrhotic livers. **d**, Violin plot: number of total Unique Molecular Identifiers (nUMI) expressed across liver-resident cell lineages in healthy versus cirrhotic livers. **e**, Pie charts: proportion of cell lineage per liver sample. **f**, Box and whisker plot: agreement in expression profiles across liver samples. Pearson correlation coefficients between average expression profiles for cell in each lineage, across all pairs of samples. Black bar, median value; box edges, 25^th^ and 75^th^ percentiles; whiskers, range. **g**, t-SNE visualisation: liver-resident dataset per liver sample.

**Extended Data Figure 3:**
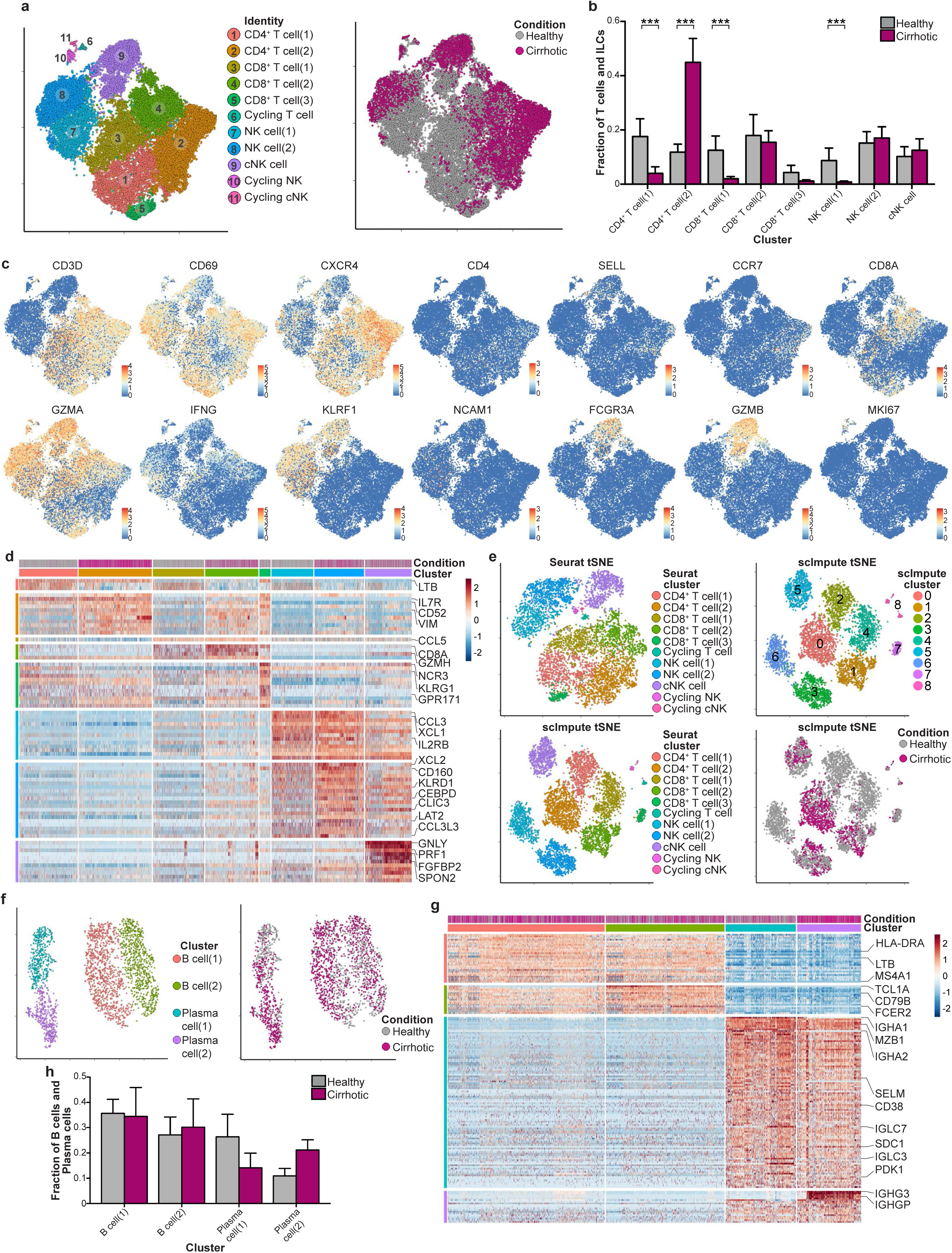
Annotating human liver lymphoid cells. **a**, t-SNE visualisation: clustering 36,900 T cells and innate lymphoid cells (ILC), annotating injury condition. cNK, cytotoxic NK cell. **b**, Fractions of T cell and ILC subpopulations in healthy (n=5) versus cirrhotic (n=5) livers, Wald. **c**, t-SNE visualisations: selected genes expressed in the T cell and ILC lineage. **d**, Scaled heatmap (red, high; blue, low): T cell and ILC cluster marker genes (colour coded top by cluster and condition), exemplar genes labelled right. Cells columns, genes rows. **e**, t-SNE visualisations: downsampled T cell and ILC dataset (7,380 cells) pre- and post-imputation; annotating data used for visualisation and clustering, inferred lineage, injury condition. No additional heterogeneity was observed following imputation. **f**, t-SNE visualisation: clustering 2,746 B cells and plasma cells, annotating injury condition. **g**, Scaled heatmap (red, high; blue, low): B cell and plasma cell cluster marker genes (colour coded top by cluster and condition), exemplar genes labelled right. Cells columns, genes rows. **h**, Fractions of B cell and plasma cell subpopulations in healthy (n=5) versus cirrhotic (n=5) livers, Wald. Error bars, s.e.m.; *** p-value < 0.001.

**Extended Data Figure 4:**
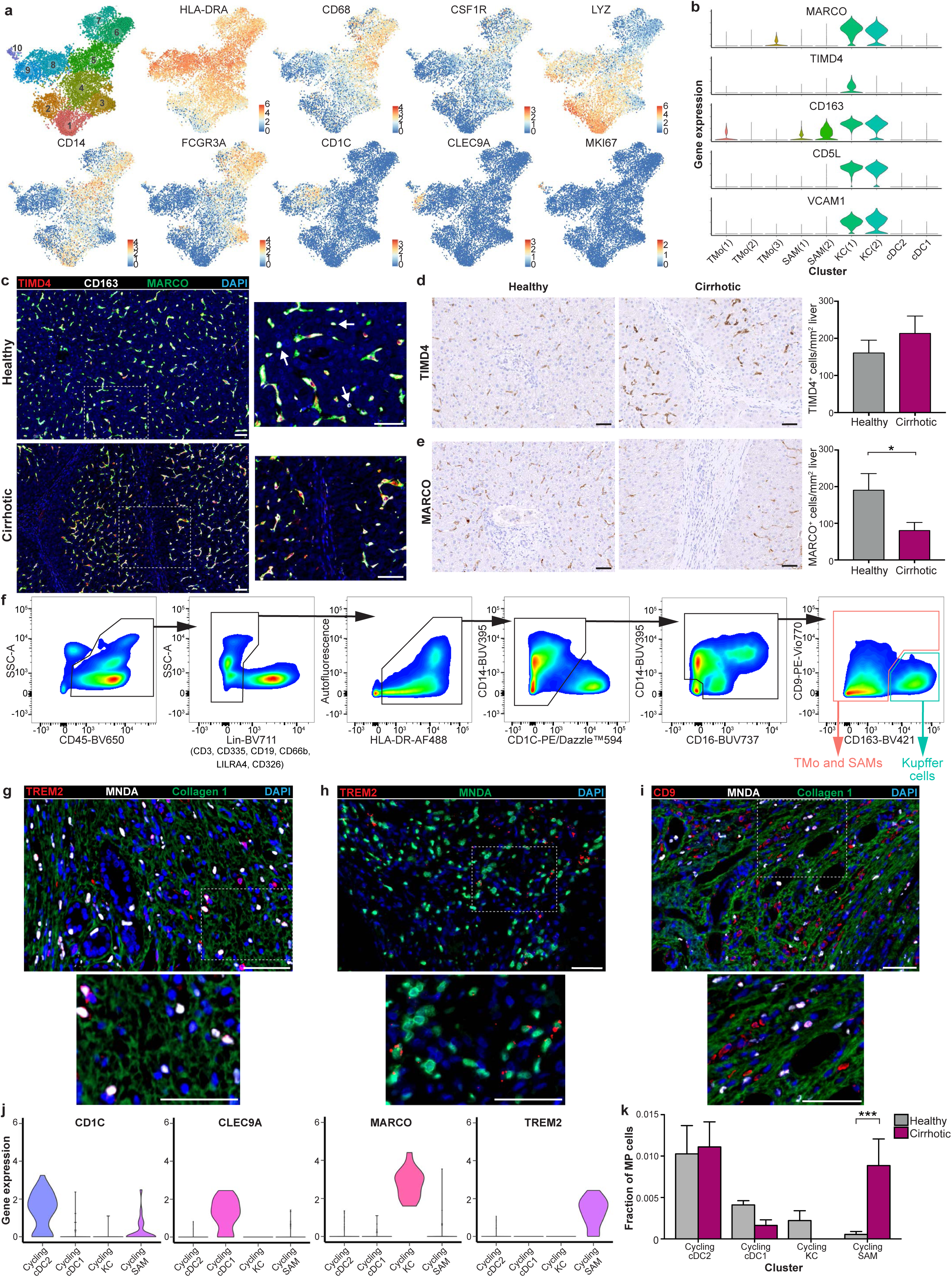
Annotating human liver mononuclear phagocytes. **a**, t-SNE visualisation: clustering and selected genes expressed in mononuclear phagocyte (MP) lineage. **b**, Violin plots: Kupffer cell (KC) cluster markers. **c**, Representative immunofluorescence micrograph, healthy versus cirrhotic liver: TIMD4 (red), CD163 (white), MARCO (green), DAPI (blue), arrows CD163^+^MARCO^+^TIMD4^-^ cells. **d**, Automated cell counting: TIMD4 staining, healthy (n=12) versus cirrhotic (n=9) liver, Mann-Whitney. **e**, Automated cell counting: MARCO staining, healthy (n=8) versus cirrhotic (n=8) liver, Mann-Whitney. **f**, Representative flow cytometry plots: gating strategy for identifying KC, TMo and SAMs. SAMs are detected as TREM2+CD9+ cells within the TMo and SAM gate (see Fig. 2f). **g**, Representative immunofluorescence micrograph, cirrhotic liver: TREM2 (red), MNDA (white), collagen 1 (green), DAPI (blue). **h**, Representative micrograph, cirrhotic liver: TREM2 (smFISH; red), MNDA (immunofluorescence; green), DAPI (blue). **i**, Representative immunofluorescence micrograph, cirrhotic liver: CD9 (red), MNDA (white), collagen 1 (green), DAPI (blue). **j**, Violin plots: cycling MP cluster markers. **k**, Fractions of cycling MP subpopulations in healthy (n=5) versus cirrhotic (n=5) livers, Wald. Scale bars, 50μm. Error bars, s.e.m.; * p-value < 0.05, *** p-value < 0.001.

**Extended Data Figure 5:**
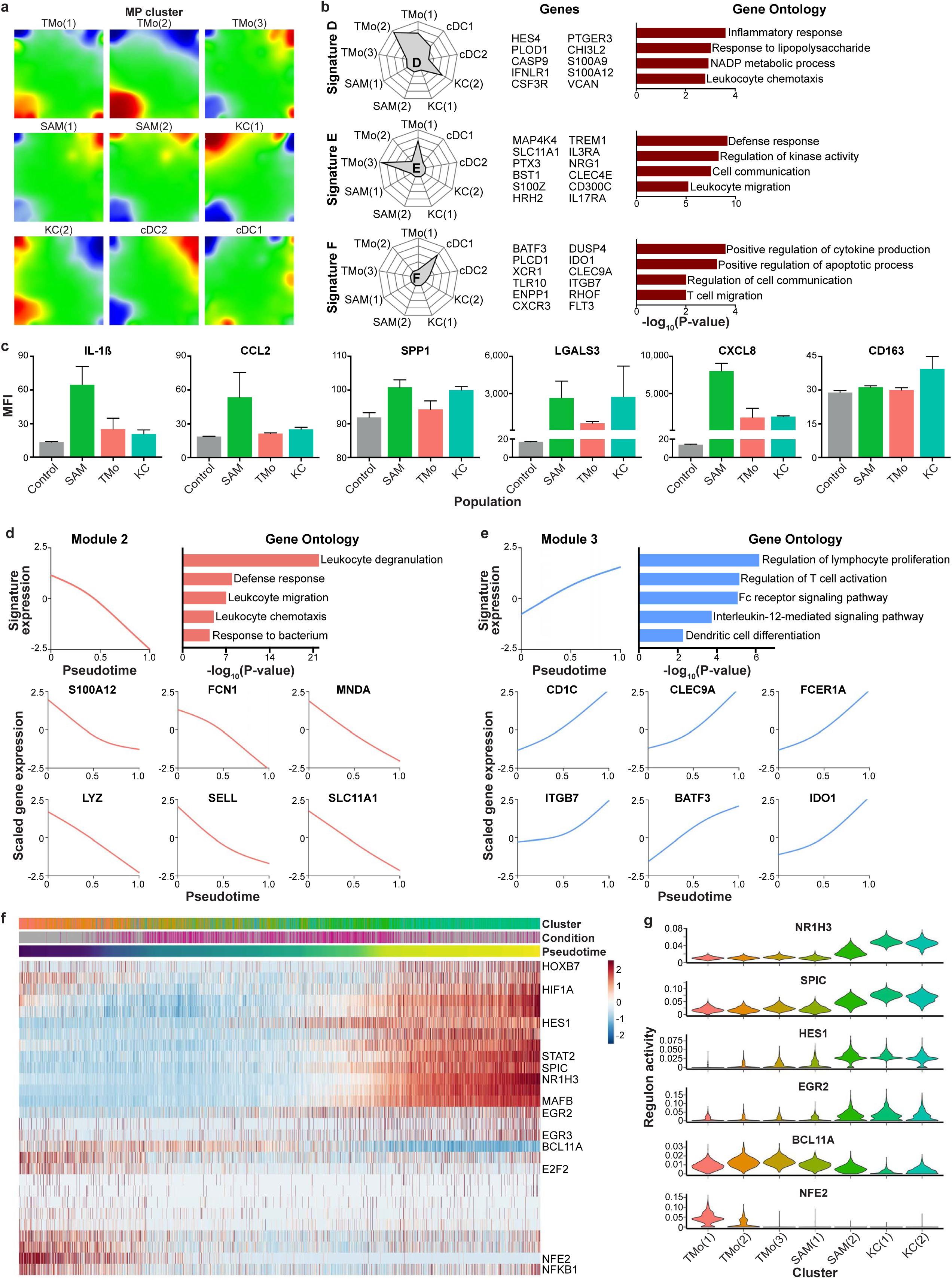
Phenotypic characterisation of mononuclear phagocytes in healthy and cirrhotic human livers. **a**, Self-Organising Map (SOM; 60×60 grid): smoothed mean metagene expression profile for mononuclear phagocyte (MP) subpopulations. **b**, Radar plots (left): metagene signatures D-F showing distribution of signature expression across MP subpopulations, exemplar genes (middle) and gene ontology (GO) enrichment (right). **c**, Luminex assay: quantification of levels of stated proteins in culture medium from FACS-isolated SAMs (n=3), KCs (n=2), TMo (n=2) and control (media alone, n=2). MFI, median fluorescence intensity. **d**, Cubic smoothing spline curve fitted to averaged expression of all genes in module 2 from blood monocyte-SAM pseudotemporal trajectory, selected GO enrichment (right) and curves fit to exemplar genes (below). **e**, Cubic smoothing spline curve fitted to averaged expression of all genes in module 3 from blood monocyte-cDC pseudotemporal trajectory, GO enrichment (right) and curves fit to exemplar genes (below). **f**, Scaled heatmap (red, high; blue, low): transcription factor regulons across MP pseudotemporal trajectory and in KCs. Colour coded top by MP cluster, condition and pseudotime, selected regulons labelled right. Cells columns, regulons rows. **g**, Violin plots: selected regulons expressed across MP clusters.

**Extended Data Figure 6:**
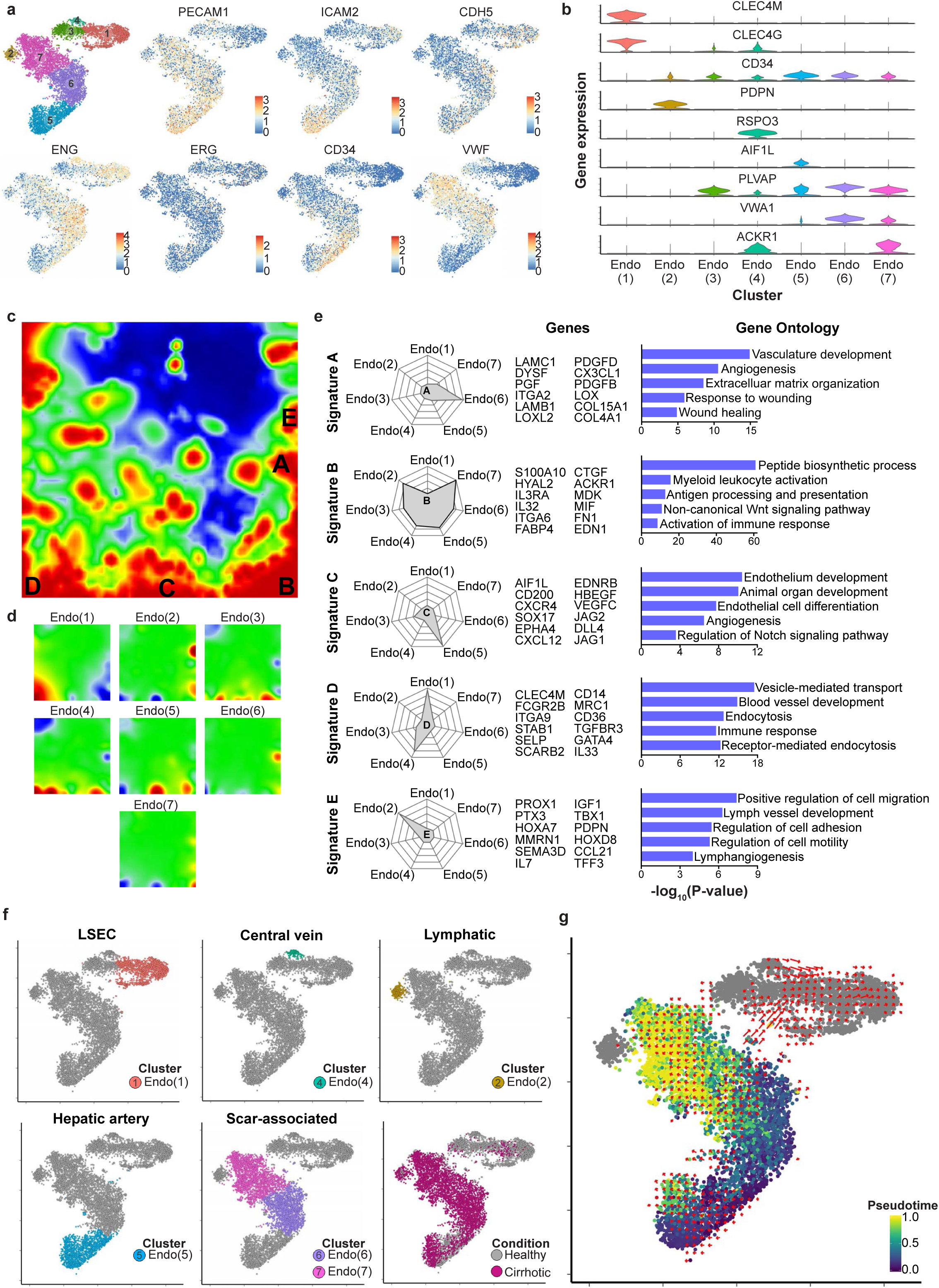
Phenotypic characterisation of endothelial cells in healthy and cirrhotic human livers. **a**, t-SNE visualisations: clusters and selected genes expressed in endothelial lineage. **b**, Violin plots: endothelial cluster marker genes. **c**, Self-Organising Map (SOM; 60 x 60 grid): smoothed scaled metagene expression of endothelia lineage. 21,237 genes, 3,600 metagenes, 45 signatures. A-E label metagene signatures overexpressed in one or more endothelial subpopulations. **d**, SOM: smoothed mean metagene expression profile for each endothelial subpopulation. **e**, Radar plots (left): metagene signatures A-E showing distribution of signature expression across endothelial subpopulations, exemplar genes (middle) and gene ontology (GO) enrichment (right). **f**, t-SNE visualisation: endothelia subpopulation annotation, injury condition. **g**, t-SNE visualisation: endothelial lineage annotated by *monocle* pseudotemporal dynamics (purple to yellow; grey indicates lack of inferred trajectory). RNA velocities (red arrows) visualised using Gaussian smoothing on regular grid.

**Extended Data Figure 7:**
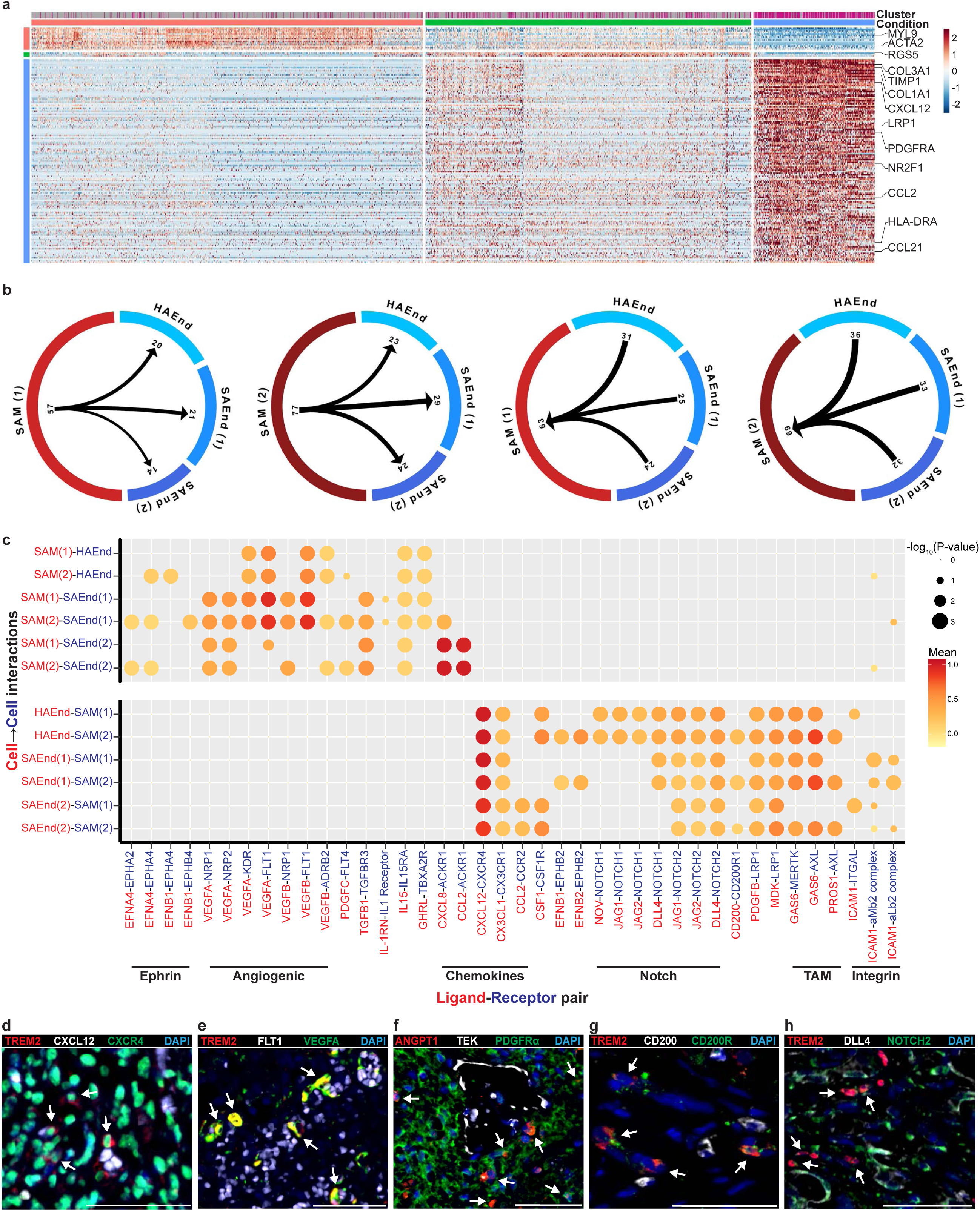
Scar-associated macrophage and endothelial cell interactions in the fibrotic niche. **a**, Scaled heatmap (red, high; blue, low): mesenchymal cluster marker genes (colour coded top by cluster and condition), exemplar genes labelled right. Cells columns, genes rows. **b**, Circle plots: potential interaction magnitude between ligands expressed by and receptors expressed on scar-associated macrophages and endothelial cells. **c**, Dotplot: selected ligand-receptor interactions between scar-associated macrophages and endothelial cells in fibrotic niche. x-axis, ligand (red) and cognate receptor (blue); y-axis, ligand-expressing cell population (red) and receptor-expressing cell population (blue). P-values indicated by circle size, means of average ligand and receptor expression levels in interacting subpopulations indicated by colour (red, high; yellow, low). TAM, TAM receptor tyrosine kinases. **d to h**, Representative immunofluorescence micrographs, fibrotic niche in cirrhotic liver. **d**, TREM2 (red), CXCL12 (white), CXCR4 (green), DAPI (blue), arrows TREM2^+^CXCR4^+^ cells. **e**, TREM2 (red), FLT1 (white), VEGFA (green), DAPI (blue), arrows TREM2^+^VEGFA^+^ cells. **f**, ANGPT1 (red), TEK(white), PDGFR*α* (green), DAPI (blue), arrows ANGPT1^+^PDGFR*α*^+^ cells. **g**, TREM2 (red), CD200 (white), CD200R (green), DAPI (blue), arrows TREM2^+^CD200R^+^ cells. **h**, TREM2 (red), DLL4 (white), NOTCH2 (green), DAPI (blue), arrows TREM2^+^NOTCH2^+^ cells. Scale bars, 50μm.

