## Supplementary Table 16 for "Resolving the fibrotic niche of human liver cirrhosis using single-cell transcriptomics"

Supplementary Table 16: Antibodies used in study

| Name | Conjugate | Host | Source | Catalogue Number | Lot Number(s) | Application(s) | Fluorescent Dilution | IHC Dilution |
|---|---|---|---|---|---|---|---|---|
| **FCM/FACS Antibodies** | | | | | | | | |
| CD1C | PEDazzle 594 | Mouse | Biolegend | 331531 | B233146 | FCM | 1:100 | - |
| CD3 | Biotin | Mouse | Biolegend | 300403 | B234471 | FCM | 1:100 | - |
| CD9 | PEVio770 | Mouse | Miltenyi | 130-103-990 | 5171128423 | FCM | 1:20 | - |
| CD14 | BUV395 | Mouse | BD Biosciences | 563562 | 6293988 | FCM | 1:50 | - |
| CD16 | BUV737 | Mouse | BD Biosciences | 564433 | 7201879 | FCM | 1:50 | - |
| CD19 | Biotin | Mouse | Biolegend | 302203 | B213806 | FCM | 1:100 | - |
| CD45 | APC | Mouse | Biolegend | 304011 | B190801 | FACS | 1:100 | - |
| CD45 | BV650 | Mouse | BD Biosciences | 563717 | 7005750 | FCM | 1:50 | - |
| CD66b | Biotin | REA | Miltenyi | 130-104-412 | 5171128425 | FCM | 1:20 | - |
| CD66b | FITC | Mouse | Biolegend | 305103 | B213990<br>B209380 | FACS | 1:200 | - |
| CD163 | BV421 | Mouse | BD Biosciences | 566277 | 7256971 | FCM | 1:20 | - |
| CD326 | Biotin | Mouse | Biolegend | 324215 | B205338 | FCM | 1:100 | - |
| CD335 | Biotin | REA | Miltenyi | 130-112-276 | 5171128399 | FCM | 1:50 | - |
| HLA-DR | AF488 | Mouse | Biolegend | 307619 | B228888 | FCM | 1:50 | - |
| LILRA4 | Biotin | REA | Miltenyi | 130-097-756 | 5171128371 | FCM | 1:11 | - |
| TREM2 | APC | Rat | R&D | FAB17291A | AAD5051591 | FCM | 1:20 | |
| **IHC/IHCF Antibodies** | | | | | | | | |
| ACKR1 | - | Rabbit | Atlas Antibodies | HPA017672 | R07143 | IHC/IHCF | 1:50 | 1:50 |
| AIF1L | - | Rabbit | Atlas Antibodies | HPA020522 | A116927 | IHCF | 1:500 | - |
| CD9 | - | Rabbit | Abcam | AB92726 | GR260186-36 | IHC/IHCF | 1:2000 | 1:4000 |
| CD34 | - | Rabbit | Abcam | AB81289 | GR305367-24 | IHCF | 1:100 | - |
| CD163 | - | Mouse | Atlas Antibodies | HPA046404 | A107257 | IHCF | 1:100 | - |
| CLEC4M | - | Rabbit | Sigma | HPA042661 | A107219 | IHC/IHCF | 1:100 | 1:200 |
| Collagen 1 | - | Rabbit | Abcam | AB34710 | GR319289-2 | IHCF | 1:300 | - |
| CXCL12 | - | Mouse | R&D Systems | MAB350 | COJ0517091 | IHCF | 1:100 | - |
| CXCR4 | - | Mouse | R&D Systems | MAB172 | AVB081805A | IHCF | 1:500 | - |
| MARCO | - | Rabbit | Atlas Antibodies | HPA063793 | D117074 | IHC/IHCF | 1:500 | 1:50 |
| MNDA | - | Rabbit | Atlas Antibodies | HPA034532 | A115510 | IHC/IHCF | 1:500 | 1:500 |
| PDGFRA | - | Goat | R&D Systems | AF-307-NA | VG0717081 | IHC/IHCF | 1:200 | 1:200 |
| PDPN | - | Rabbit | Atlas Antibodies | HPA007534 | F115991 | IHCF | 1:200 | - |
| PLVAP | - | Rabbit | Novus | NBP1-83911 | B115774 | IHC/IHCF | 1:200 | 1:25 |
| RSPO3 | - | Rabbit | Novus | NBP1-82234 | B105245 | IHCF | 1:500 | - |
| TIMD4 | - | Rabbit | Sigma | HPA015625 | A113789 | IHC/IHCF | 1:500 | 1:500 |
| TREM2 | - | Rabbit | Proteintech | 13483-1-AP | 0.00005004 | IHC/IHCF | 1:250 | 1:500 |
| PDGFB | - | Rabbit | Abcam | ab23914 | GR3197857-1 | IHCF | 1:100 | - |
| IL1R1 | - | Rabbit | Abcam | ab106278 | GR3207174-5 | IHCF | 1:500 | - |
| IL-1β | - | Rabbit | Abcam | ab9722 | GR309542-13 | IHCF | 1:100 | - |
| TNFRSF12A | - | Rabbit | Abcam | ab109365 | GR3218010-2 | IHCF | 1:100 | - |
| TNFSF12 | - | Goat | R&D | AF1090-SP | HKQ0115091 | IHCF | 1:100 | - |

| | | | | | | | | |
|---|---|---|---|---|---|---|---|---|
| CCL2 | - | Mouse | Novus | NBP2-2211555 | A-6 | IHCF | 1:100 | - |
| CCR2 | - | Mouse | R&D | MAB150-SP | AOT0218041 | IHCF | 1:100 | - |
| Notch2 | - | Rabbit | Atlas Antibodies | HPA048743 | R58685 | IHCF | 1:500 | - |
| Notch3 | - | Rabbit | Abcam | ab23426 | GR3226374-1 | IHCF | 1:500 | - |
| Dll4 | - | Rabbit | Abcam | ab7280 | GR211685-45 | IHCF | 1:500 | - |
| CXCL12 | - | Mouse | R&D | MAB350-SP | C0J0517091 | IHCF | 1:100 | - |
| CXCR4 | - | Mouse | R&D | MAB172-SP | AVB1118051 | IHCF | 1:500 | - |
| FLT1 | - | Rabbit | Abcam | ab32152 | GR157619-54 | IHCF | 1:500 | - |
| VEGFA | - | Rabbit | Abcam | ab52917 | GR3219705-1 | IHCF | 1:250 | - |
| ANGPT1 | - | Goat | R&D | AF923-SP | GPN1117011 | IHCF | 1:200 | - |
| TEK | - | Goat | R&D | AF313-SP | CMF0518031 | IHCF | 1:100 | - |
| CD200 | - | Goat | R&D | AF2724-SP | VJZ021806A | IHCF | 1:100 | - |
| CD200R | - | Rabbit | Bioss/Insight | bs-109SR | 980698W | IHCF | 1:200 | - |

FCM: Flow cytometry; FACS: Fluorescence-activated cell sorting; IHC: Immunohistochemistry; IHCF: Fluorescent immunohistochemistry; REA: REAffiinity
